# Emergence of distinct and heterogeneous strains of amyloid beta as Alzheimer’s disease progresses in Down syndrome

**DOI:** 10.1101/2021.07.30.454527

**Authors:** Alison M. Maxwell, Peng Yuan, Brianna M. Rivera, Wilder Schaaf, Mihovil Mladinov, Vee P. Prasher, Andrew C. Robinson, William F. DeGrado, Carlo Condello

**Affiliations:** Department of Pharmaceutical Chemistry, Cardiovascular Research Institute, University of California, San Francisco, CA 94158; Institute for Neurodegenerative Diseases, Weill Institute for Neurosciences, University of California, San Francisco, CA 94158; Department of Physics & Astronomy, San Francisco State University, San Francisco, CA 94132; Memory & Aging Center, Weill Institute for Neurosciences, University of California, San Francisco, CA 94158; South Birmingham Community NHS Trust, Birmingham, UK; Liverpool John Moores University, Liverpool, UK; Division of Neuroscience & Experimental Psychology, Faculty of Biology, Medicine and Health, School of Biological Sciences, The University of Manchester, Salford Royal Hospital, Salford, UK; Department of Neurology, Weill Institute for Neurosciences, University of California, San Francisco, CA 94158

## Abstract

Amyloid beta (Aβ) is thought to play a critical role in the pathogenesis of Alzheimer’s disease (AD). Prion-like Aβ polymorphs, or “strains”, can have varying pathogenicity and may underlie the phenotypic heterogeneity of the disease. In order to develop effective AD therapies, it is critical to identify the strains of Aβ that might arise prior to the onset of clinical symptoms and understand how they may change with progressing disease. Down syndrome (DS), as the most common genetic cause of AD, presents promising opportunities to compare such features between early and advanced AD. In this work, we evaluate the neuropathology and Aβ strain profile in the post-mortem brain tissues of 210 DS, AD, and control individuals. We assayed the levels of various Aβ and tau species and used conformation-sensitive fluorescent probes to detect differences in Aβ strains among individuals and populations. We found that these cohorts have some common but also some distinct strains from one another, with the most heterogeneous populations of Aβ emerging in subjects with high levels of AD pathology. The emergence of distinct strains in DS at these later stages of disease suggests that the confluence of aging, pathology, and other DS-linked factors may favor conditions that generate strains that are unique from sAD.

## 1. INTRODUCTION

Alzheimer’s disease (AD) is a progressive neurodegenerative disease that affects 10% of the US population over 65 years of age^1^ and responds only minimally to currently available therapeutics.^2^ Most people with AD initially suffer from memory loss, apathy, and depression, followed by impaired communication and confusion, and eventually, motor debilitations that often lead to death.^3^ Inherited or familial AD (fAD) is early-onset but relatively rare, while the majority of cases are either associated with Down syndrome (AD-DS) or are sporadic (sAD), which is manifested by later onset and no clear genetic cause.

The AD brain is marked by an abundance of extracellular amyloid beta (Aβ)-rich plaques and intraneuronal neurofibrillary tangles (NFTs) composed of hyperphosphorylated tau. Both Aβ and tau are therefore believed to play a causative role in AD. However, the precise roles of these peptides—either independent or synergistic^4^—in its pathogenesis are unclear. Aβ aggregation is speculated to be the nucleating event in AD, as it accumulates in the brain 10-20 years before the onset of dementia,^5,6^ followed by tau deposition concomitant with clinical symptoms.^7–9^ The molecular genetics of AD further highlights the importance of Aβ in disease pathogenesis: mutations in the amyloid precursor protein (APP, from which Aβ is generated) or in the APP processing enzyme presenilin lead to fAD.^10^ Alternatively, mutations in tau lead to other types of dementias with NFT pathology.^11^

The onset, progression, and severity of symptoms in sAD are diverse.^12^ This heterogeneity is likely due at least in part to the structural diversity of Aβ species in the AD brain.^13–15^ Normal APP processing results in Aβ peptides of various lengths, most commonly comprising Aβ residues 1-40 (Aβ40) and 1-42 (Aβ42). These isoforms can in turn adopt a multitude of distinct molecular conformations *in vitro*,^16^ form fibrils of differing structure and pathogenicity,^17^ and have been found as diverse ultrastructural assemblies in different clinical AD phenotypes.^18,19^ The ability of brain-derived Aβ fibrils to propagate their structure in a prion-like mechanism^14,20–22^ suggests that structurally distinct, self-propagating strains of Aβ might underlie the clinicopathological heterogeneity in sAD. Indeed, we recently showed that different Aβ strains differentiate plaques in fAD subtypes,^14^ supporting a hypothesis that individuals each have only a few of the many Aβ strains found across AD. Together, this evidence suggests that different molecular structures of Aβ have varying pathogenicity and may underlie the phenotypic heterogeneity of sAD. Understanding whether more pathogenic strains are seeded early in the disease or evolve with time and environmental changes would enable more targeted approaches to diagnostic and therapies.

Thus, robust methods of interrogating the role of Aβ early in AD are needed. Previous efforts have focused on fAD individuals with mutations that affect the production of Aβ because they cause comparable phenotypes to sAD in an identifiable, deterministic manner.^23^ Such studies have yielded critical insights into the pathogenesis of AD, but as fAD represents <1% of all AD cases,^23^ the size and scope of these investigations are limited. Alternatively, as the most common genetic cause of AD, Down syndrome (DS) presents promising opportunities to study the onset of AD. Due to trisomy of chromosome 21 (Chr21), which encodes *APP*,^24^ people with DS have a lifelong overproduction of APP leading to increased accumulation of Aβ. AD neuropathology is prevalent in DS individuals over the age of 40,^25^ while dementia is diagnosed in approximately 65-80% of the DS population over 65 years of age.^26^ The distribution and biochemical composition of Aβ plaques and NFTs in AD-DS are similar to fAD and sAD,^27,28^ as is the progression of clinical symptoms including dementia.^29,30^ Thus, compared to the relatively rare fAD, DS offers unique advantages for comprehensive studies of AD pathogenesis.

Despite promising prospects of AD prevention trials in DS,^31^ research into the molecular pathogenesis of AD-DS has been limited by a number of obstacles. A lack of standardized collection and documentation for DS autopsy cases has restricted the size and characterization of study cohorts.^32^ Furthermore, due to the overexpression of APP and other Chr21 genes in AD-DS, its molecular phenotypes and mechanisms may be different from sAD. Yet the histological methods often used for assessing the distribution and morphology of Aβ and tau lesions in DS have often lacked the specificity to interrogate such molecular detail. PET imaging, while enabling longitudinal studies of the spread and severity of Aβ load in AD-DS, is also relatively nonspecific to Aβ morphotypes^33–35^ and primarily binds only a subfraction of Aβ in AD.^36^ Clearly, there is a need for applying precise, high-resolution methods to the analysis of Aβ pathology in AD-DS.

Environment-sensitive fluorescent dyes such as Congo Red,^37^ ThT,^38^ and others^39^ have historically been invaluable in probing Aβ conformation in AD. Though lacking the definitive structural detail of cryo-electron microscopy (cryo-EM) or solid-state nuclear magnetic resonance (ssNMR), dye-based analysis is high throughput while still sensitive to structural differences.^40^ Of further advantage, it can be performed *in situ*, without the need for stringent purification. We previously optimized a set of three dyes, BF-188, FSB, and curcumin, to discriminate amyloid deposits in post-mortem fAD and sAD tissue and identified distinct Aβ strains within individuals.^14^ Because differences in fibril conformation, isoform composition, density, and other local environmental factors can all impact a plaque’s fluorescence signature, they contribute to the definition of different strains in this context.

Here we apply this method in the first comparative analysis of Aβ strains among plaques in 210 individuals with DS or AD as well as in control subjects. We sought to identify whether a distinct subset of Aβ strains are present before the onset of dementia in AD and how such strains might change or persist throughout the disease. Using principal component analysis (PCA) on the fluorescence spectra of dyes bound to intact plaques, we found that most strains of Aβ appear to be common to sAD and AD-DS. However, some DS individuals with the most severe neuropathology additionally present with some distinct strains. These differences are partially but not fully explained by the bulk amount of Aβ40 and Aβ42 in each tissue and may be related to a 2-fold elevation of phospho-Tau (pTau) in our AD-DS cohort. We posit that the increasingly divergent biochemical environment of the aging DS brain may be able to foster the propagation of unique strains of Aβ not otherwise found in AD.

## 2. RESULTS

### 2.1. AD pathology varies by age and cohort

We first sought to characterize the key biological and genetic attributes of the cases in our study to allow us to later control for potential covariates in our analysis of Aβ strains. The age distributions of the three main cohorts—DS (including AD-DS), AD (without DS), and ADNC—are shown in **Figure 1A.** We prioritized obtaining and analyzing DS cases under 40 years of age, since these were the most likely to provide insight into pre-clinical AD. Though such cases are relatively rare, we obtained 21 cases between 20-40 years of age, making this the largest study of young DS post-mortem tissue to date in addition to the largest known cohort of DS generally in a molecular study of Aβ. The majority of our DS cases were aged 35-65 years at the time of death, while most AD cases tended to be older (aged ~55-90 years); generally, these two groups are considered comparable because of accelerated aging in DS. Fifty percent were male and 50% female (**Figure 1C**).

**Figure 1.**
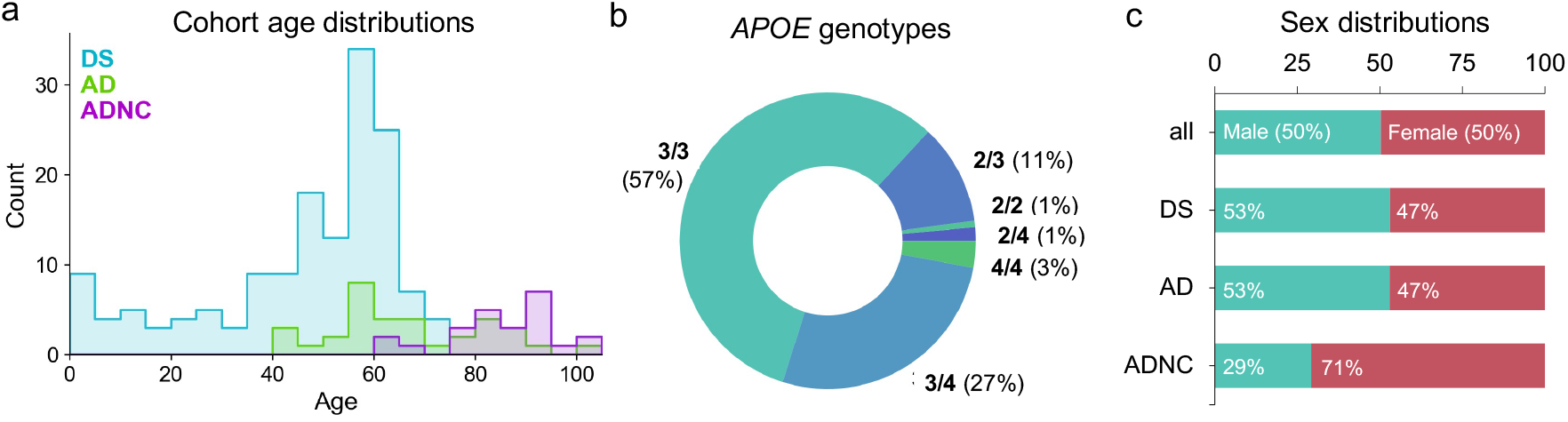
A broad range of case ages, genotypes, and sexes comprise the study cohorts. (**A**) Per-cohort distributions of patient ages at death (nDS=152, nAD=34, nADNC = 24) (**B**) Proportion of cases with each *APOE* genotype, when known (n=137). Genotypes were established through Sanger sequencing of SNP-containing *APOE* amplicons purified from gDNA. (**C**) Percent of all cases and of each cohort that were known to be male or female (n=204).

We chose to genotype our samples for *APOE*, a gene encoding the cholesterol transport and lipid metabolism protein apolipoprotein E (apoE), because its three isoforms (ε2, ε3, and ε4) are linked to varying AD severity.^41^ The ε4 allele is strongly associated with earlier onset of dementia in both DS^42^ and the general population.^43^ Genotyping revealed a majority of cases as *APOE* 3/3, with 43 cases having at least one ε4 allele (**Figure 1B**). For the subset of cases for which *APOE* was provided by the tissue bank, 90% of genotypes matched those established by our method (data not shown).

An additional source of variation in our cohorts stems from the fact that the tissue for this study was sourced from eleven different brain banks and spanned nearly four decades of collection (see **Table 1**, attached). As a consequence, the methods and timing of tissue fixation and storage post-mortem, as well as the methods and quantity of clinical analyses and neuropathological assessment at autopsy, varied greatly across our 210 samples. To obtain a standardized measure of AD neuropathology, we generated our own pathological scores based on Aβ (*X*_Aβ_) and tau (*X*_tau_) load in the frontal cortex detected using antibodies targeting Aβ40, Aβ42, and phosphorylated tau, as outlined in **Table 2**. Examples of the appearance of Aβ and tau pathology in cases assigned *X*_Aβ_ and *X*_tau_ 1 and 4 are shown in **Figure 2A**. We validated our approach by comparing our scores to the limited Braak and CERAD data that was available and found our metrics to be consistent (**Figure 2—figure supplement 1B**).

**Table 1.**
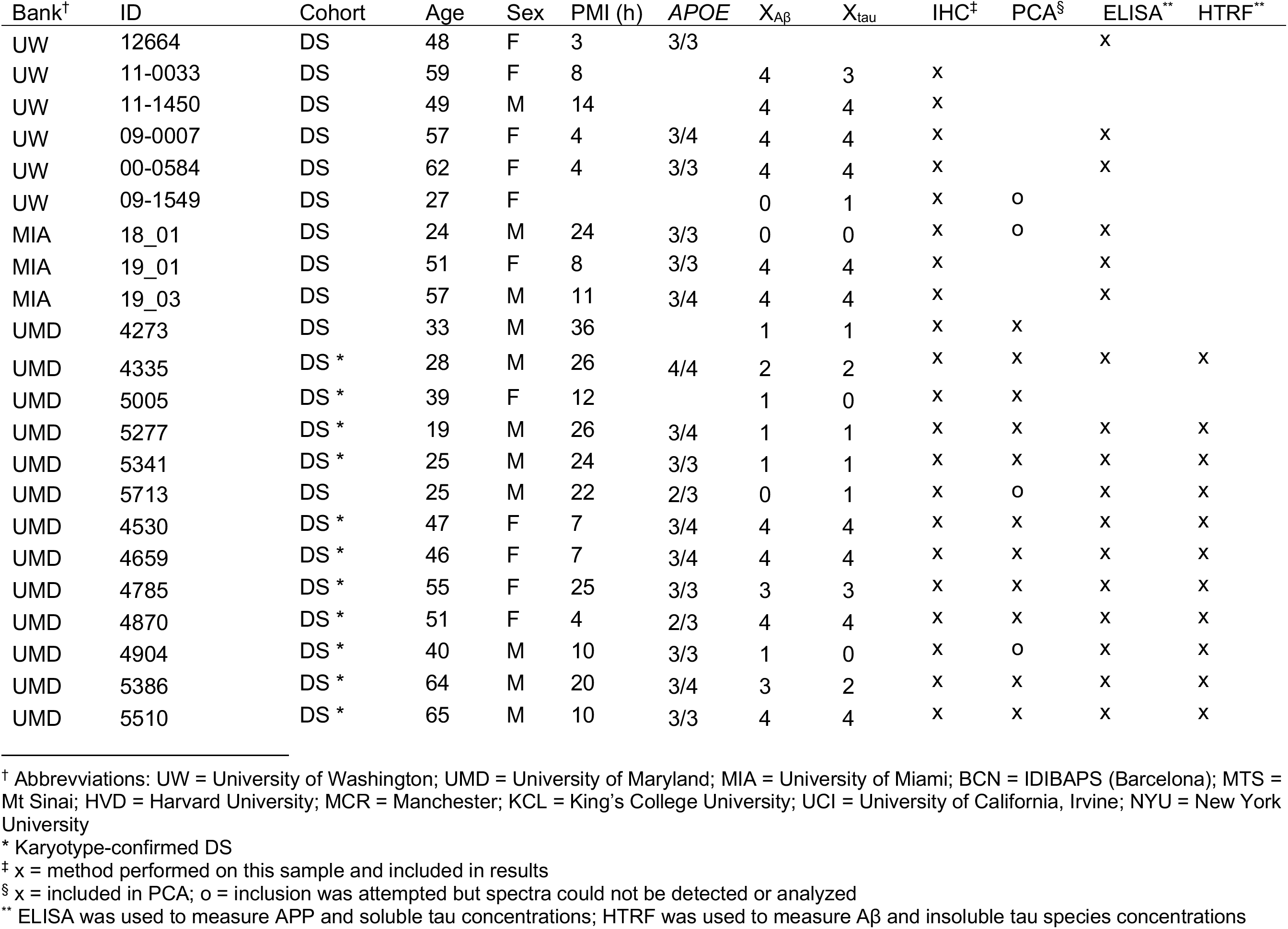
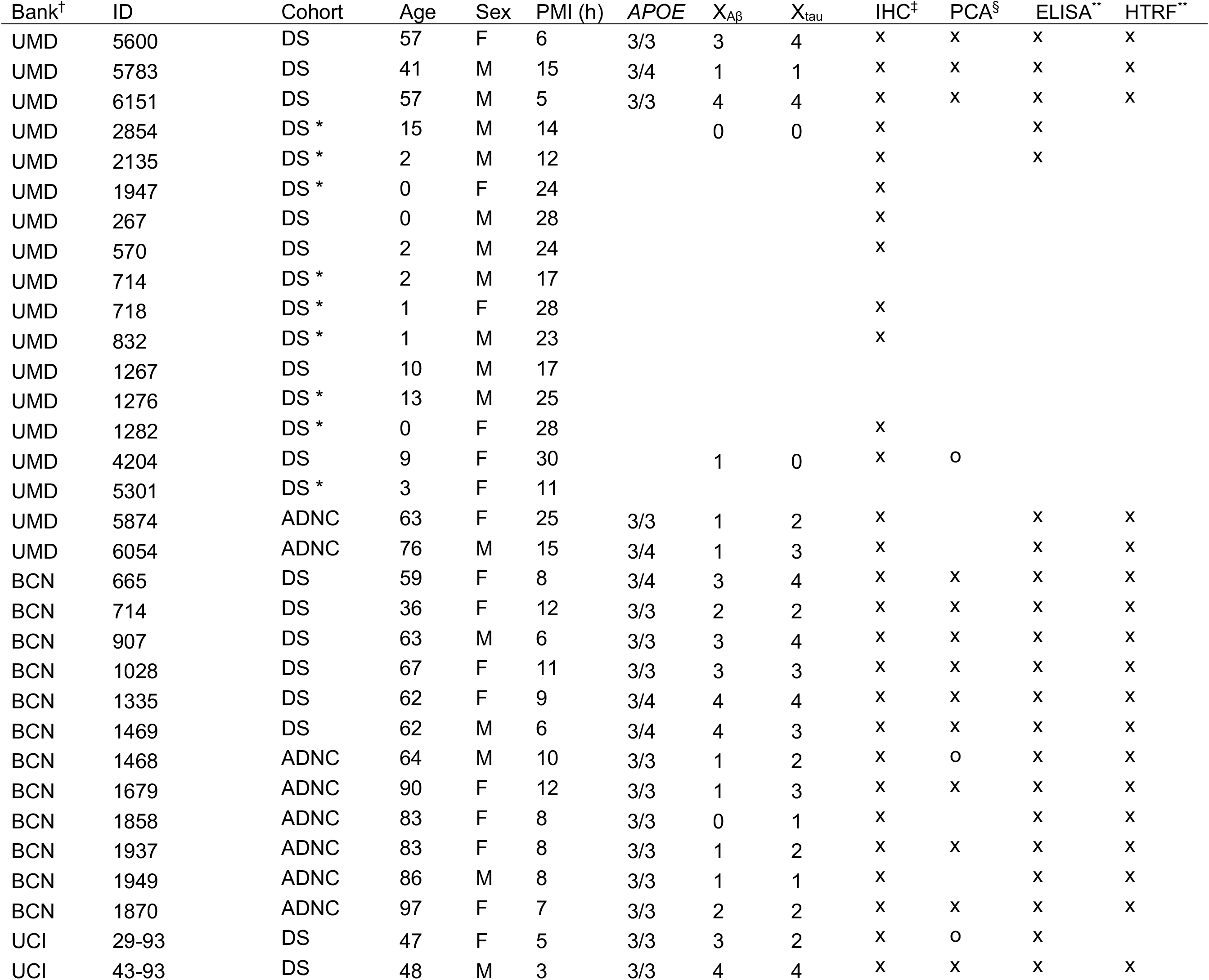
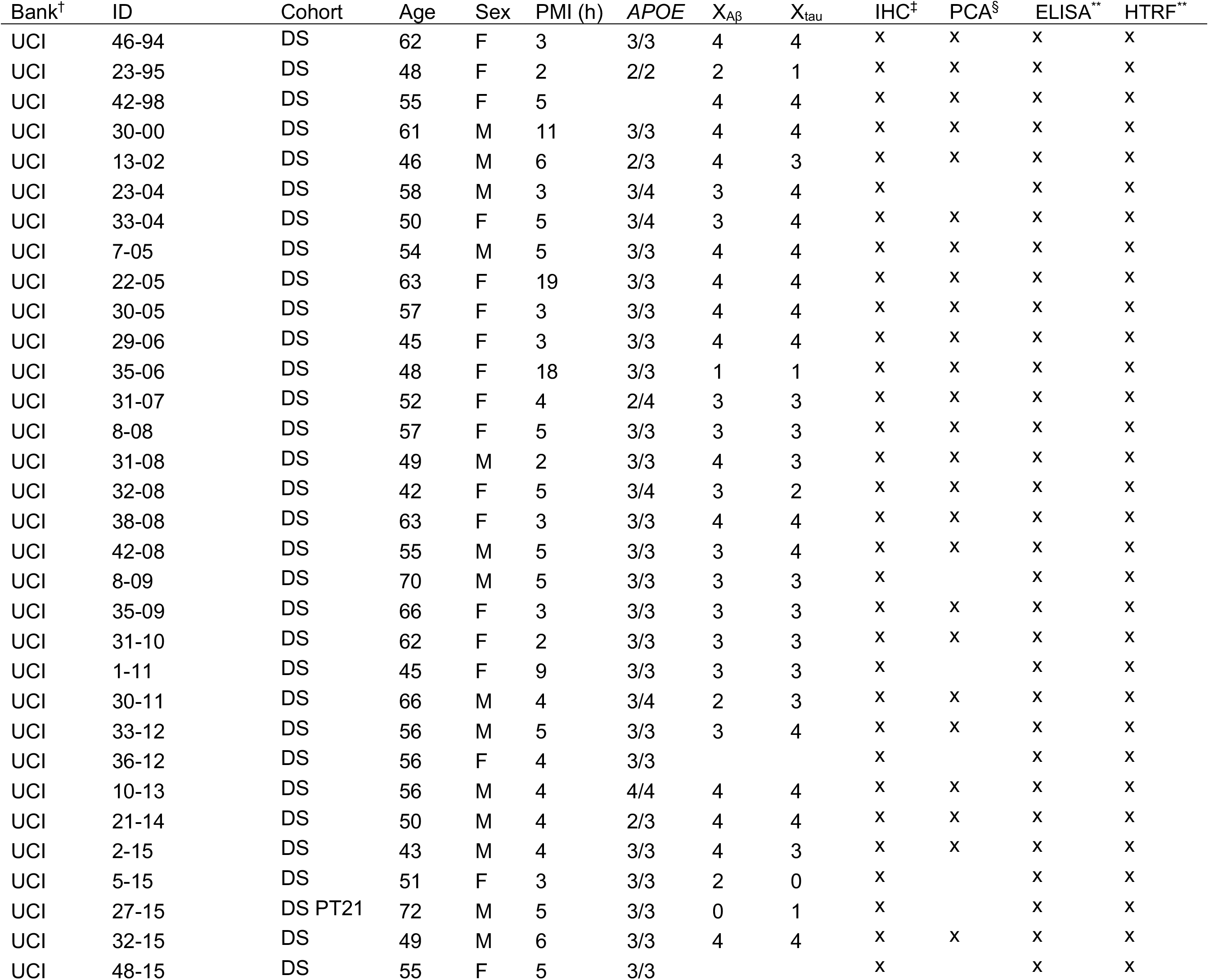
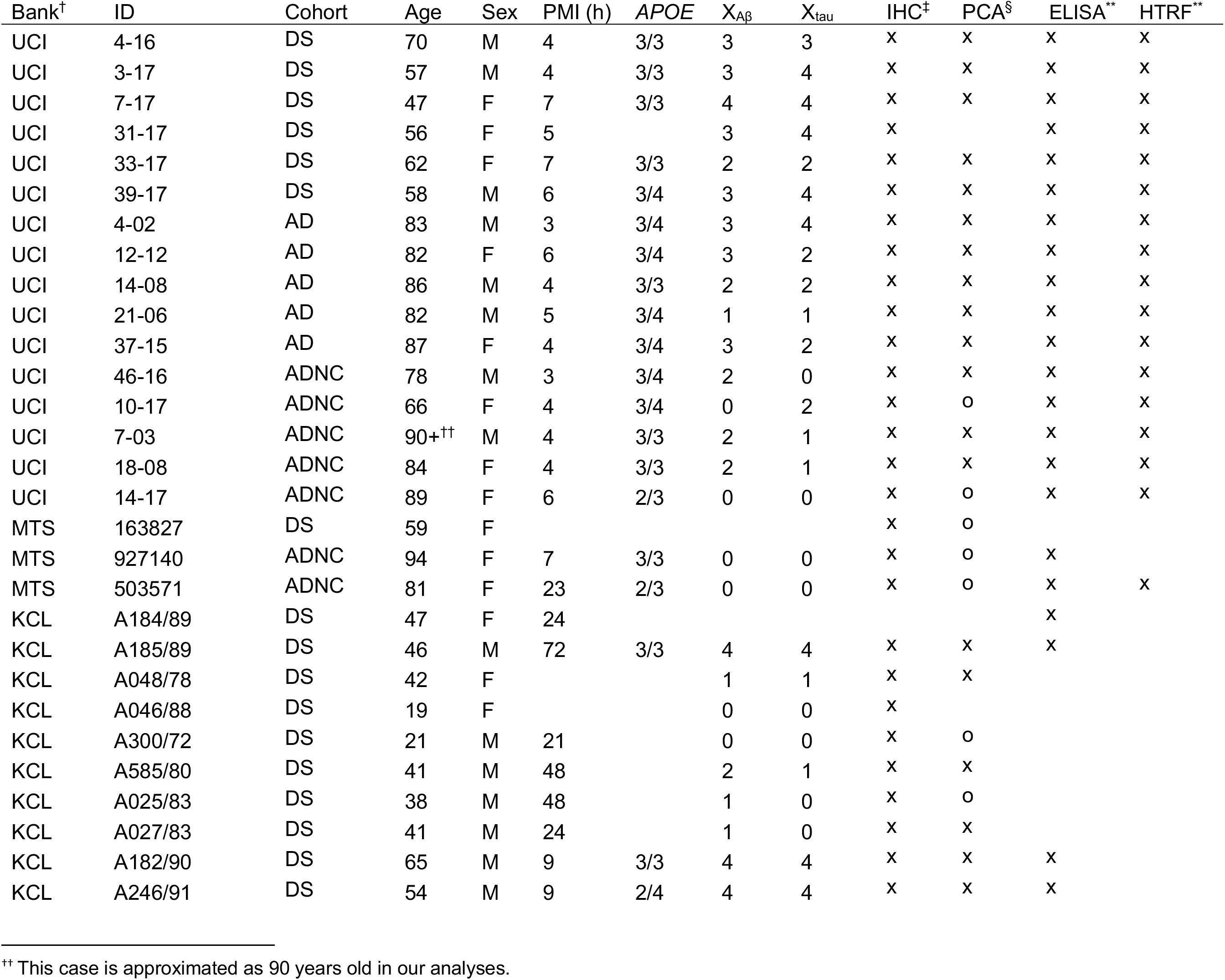
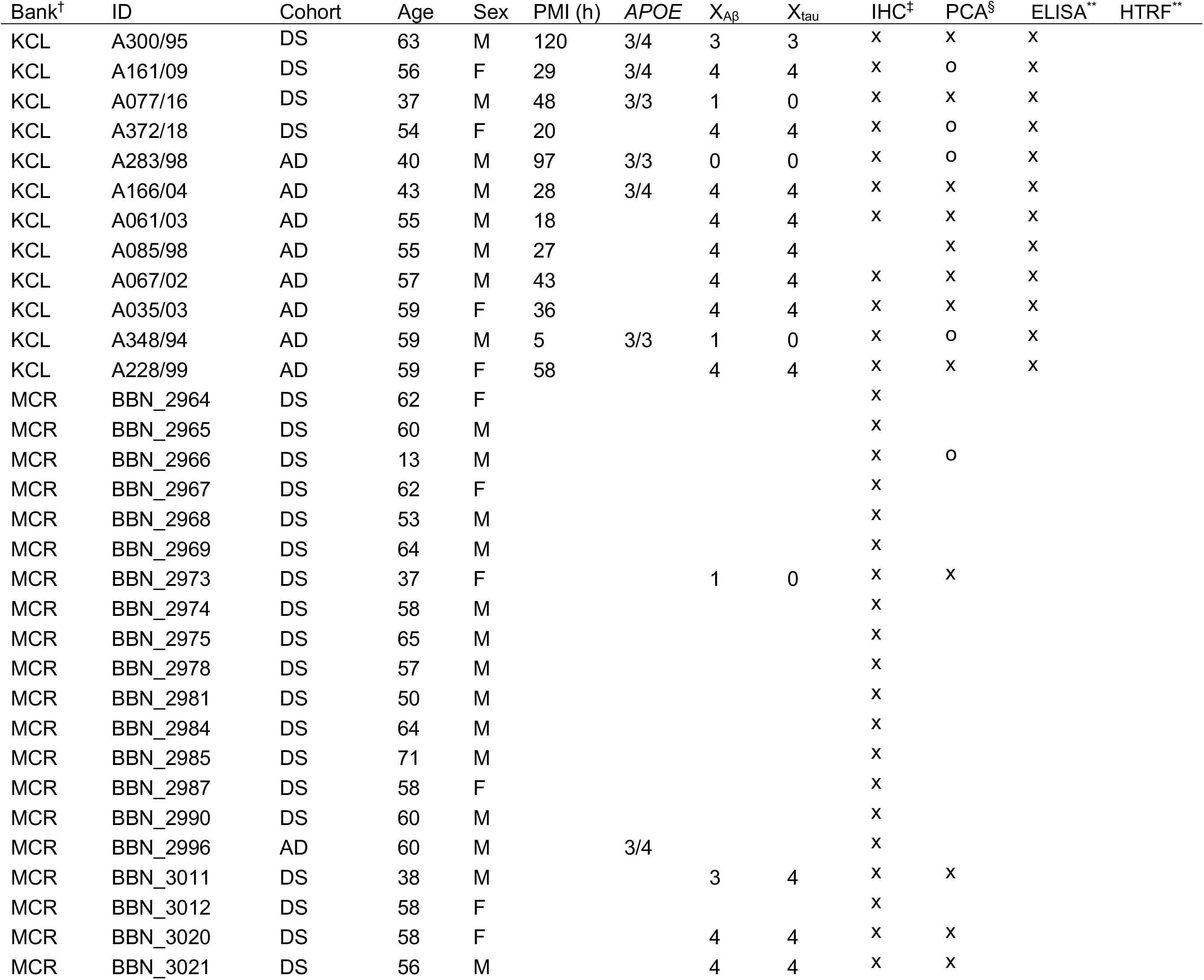
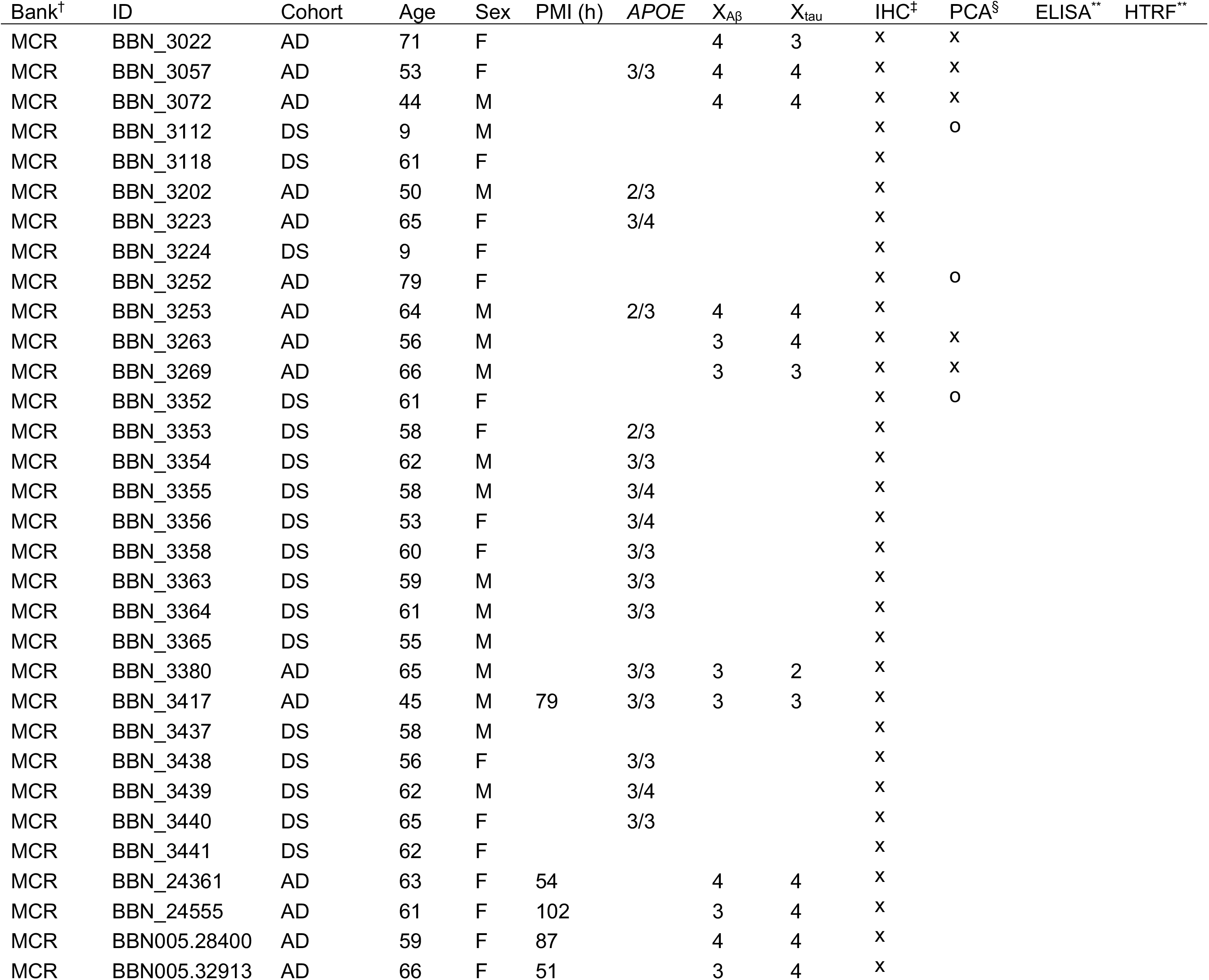
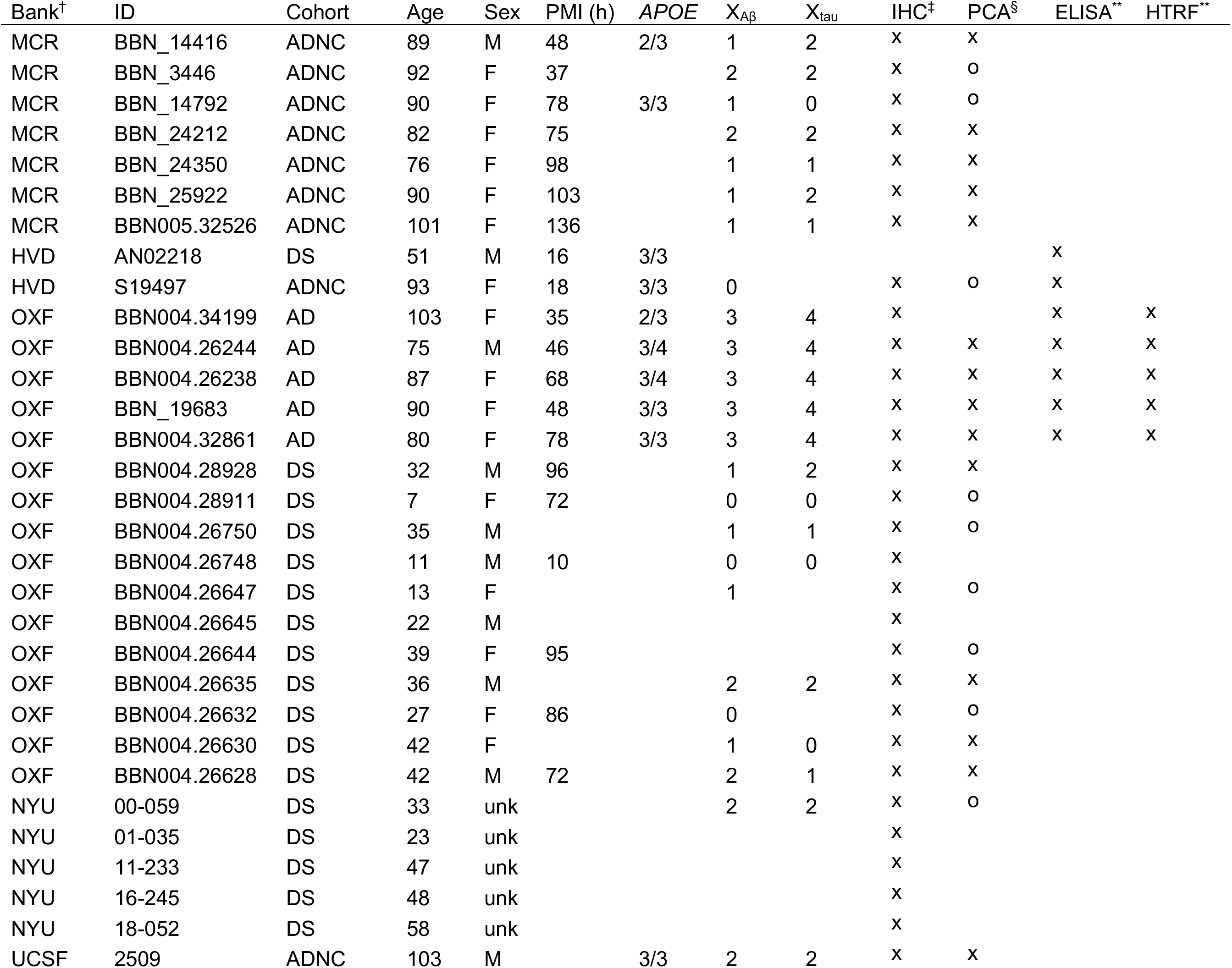
Biological data and experimental inclusion of each case.

**Table 2.** Aβ and tau scoring criteria by number of pathological markers per mm^2^

| $X$ | Aβ | Tau |
| --- | --- | --- |
| 0 | <1 plaque/mm<sup>2</sup> | <1 mature NFT/mm<sup>2</sup> |
| 1 | <1 dense-cored plaque but $\geq 2$ total plaques | 1-5 NFTs |
| 2 | $\geq 1$ dense-cored but <2 neuritic plaques | 5-12 NFTs |
| 3 | $\geq 5$ dense-cored and 2-15 neuritic plaques | 12-25 NFTs |
| 4 | $\geq 15$ neuritic plaques | $\geq 25$ mature NFTs |

**Figure 2.**
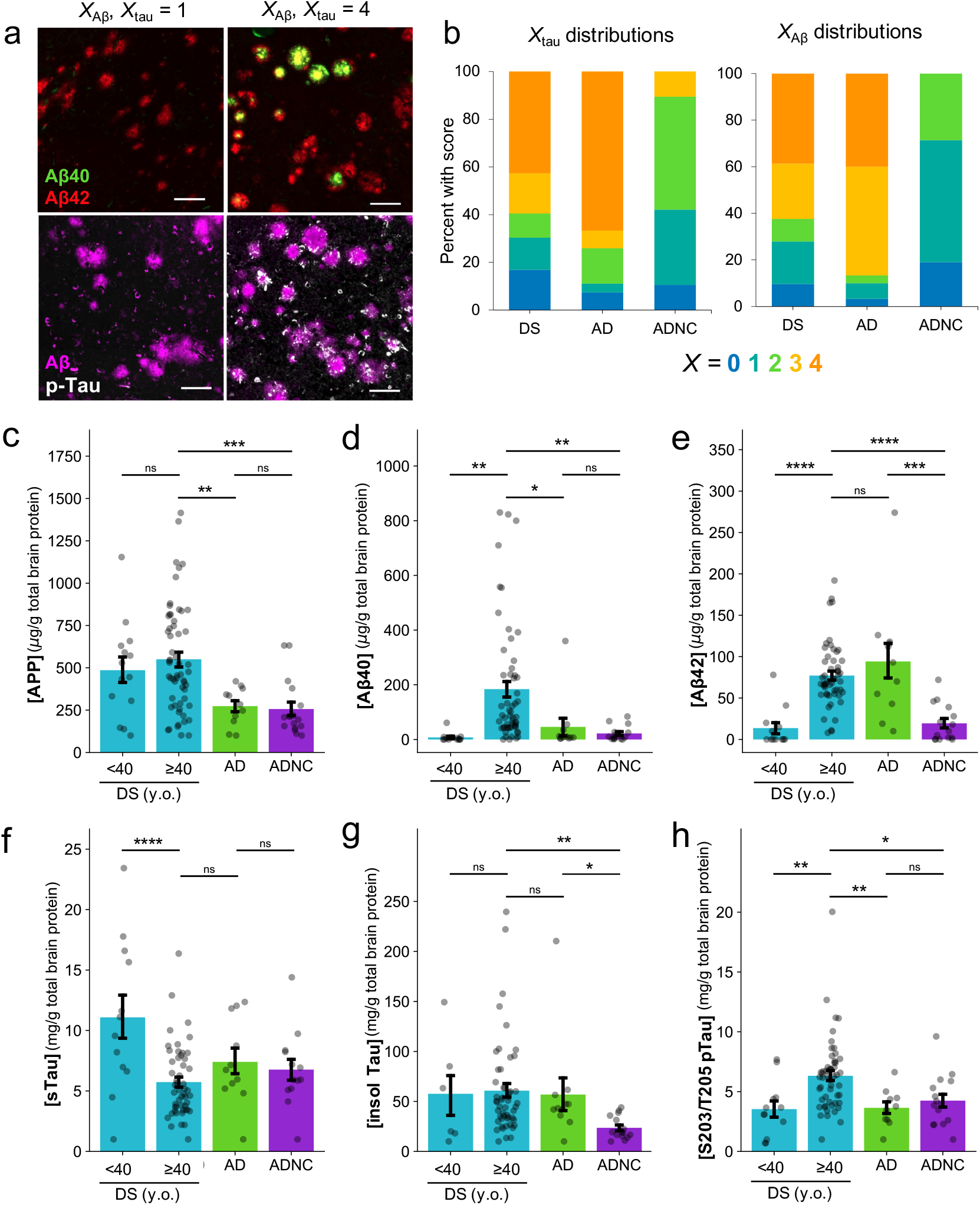
Characterization of neuropathology using custom histological scoring and biochemistry. (**A**) Representative IHC images from a DS case with X_tau_ = 1 and X_Aβ_ = 1 (UCI 35-06) and from a DS cases with X_tau_ = 4 and X_Aβ_ = 4 (UCI 29-06). FFPE sections were dual stained with primary antibodies specific for either Aβ40/Aβ42 or total Aβ/S262 pTau and were detected using fluorescent secondary antibodies. Scale bars are 100 μm. (**B**) Proportions of cases in each cohort with each *X*_Aβ_ and *X*_tau_ as determined by custom manual scoring methodology. (**C-H**) Protein concentrations determined in frozen tissue, ± SEM. For soluble proteins (APP, sTau), clarified brain homogenate was assayed by ELISA. For insoluble proteins (Aβ40, Aβ42, total insoluble tau, and pTau), formic acid-extracted samples were assayed by HTRF. Significance values were determined by student’s two-tailed t-test. *: 0.01 < p ≤ 0.05; **: 0.001 < p ≤ 0.01, ***: 0.0001 < p ≤ 0.001.

Our standardized method allows for direct, consistent comparisons of neuropathology among cases for this study. It is important to note that the absence of tau in the particular tissue section we studied does not preclude its presence elsewhere in the brain, nor are we diagnosing AD with this method. However, because tau accumulation correlates with the onset of dementia,^8,9,44^ we interpret higher *X*_tau_ to correspond to a greater likelihood of having had clinical symptoms of AD at the time of death. Specifically, we considered *X*_tau_ to indicate subjects who more likely did not experience clinical symptoms of AD during life (*X*_tau_=0) and those who likely did (*X*_tau_=4), which was generally supported by Braak staging when available (**Figure 2—figure supplement 1B**). This strategy enabled downstream comparisons of Aβ strains in DS versus probable AD-DS.

The proportions of cases with each Aβ and tau score are shown in **Figure 2B.** The majority of DS and AD cases were *X*_Aβ_ and *X*_tau_ ≥3, corresponding to the presence of many cored and neuritic plaques and NFTs, whereas ADNC cases tended to have less pathology (*X*_tau_ ≤2). Within DS, both *X*_Aβ_ and *X*_tau_ tended to increase with age (R^2^X_AB_=0.61, R^2^X_tau_=0.52, p<0.001), with the earliest signs of Aβ pathology visible in a 9-year-old with DS and of tau pathology in a 19-year-old with DS (**Figure 2—figure supplement 1A**). Importantly, 9 DS cases had *X*_tau_=0 with *X*_Aβ_≥1. Being the most likely to be associated with pre-clinical AD, the plaques in these cases were prioritized for later Aβ strain assessment. The oldest DS cases without any sign of Aβ or tau pathology were aged 27 and 51 years, respectively. No significant trends in *X*_Aβ_ and *X*_tau_ were observed relative to age among AD or ADNC cases.

### 2.2. Concentrations of APP and some Aβ and tau species differ among cohorts

We sought to characterize the amount of soluble APP in each sample in order to better understand how the processing of the protein might differ with age in DS. We also analyzed the amounts of various Aβ and tau peptides to bolster our neuropathological cohort comparisons and to assess potential novel trends in DS. The concentrations of APP, Aβ40, Aβ42, soluble tau, total tau, and S202/T205 pTau determined by ELISA and HTRF are shown in **Figure 2C–H.** On average, APP in DS was 2x higher than in AD or ADNC cases (**Figure 2C**, p<0.01 and p<0.0006 by student’s two-tailed t-test), which was unsurprising given its overexpression in DS. On average, Aβ40 and Aβ42 were 10x and 3x higher in DS individuals over 40 years of age compared to younger individuals (**Figure 2D–E**). By IHC, we determined that many cases with the highest Aβ40 levels also had significant vascular Aβ40 due to cerebral amyloid angiopathy (CAA, data not shown). However, the overall 4x elevation of Aβ40 in DS individuals over 40 years of age compared to AD was not consistently explained by CAA, suggesting altered APP processing favoring Aβ40 or shifted targeting of Aβ40 to plaques in AD-DS.

Soluble tau concentrations were highest in the very youngest DS cases (0-2 years of age) and steadily decreased with patient age until after age 30 (R^2^=0.42, p<0.0001; **Figure 2—figure supplement 2C**), but on average were not significantly different to those in AD or ADNC (**Figure 2F**). Total insoluble tau was significantly lower in ADNC than in DS, as we expected from those cases in which neuropathology was generally less severe by IHC. However, insoluble phospho-tau species have been shown to be one of the strongest predictors of disease severity in sAD and fAD.^4,45^ We found that pTau was only significantly higher than in either AD or ADNC in DS subjects over 40 years of age (**Figure 2H** and **Figure 2—figure supplement 2B**), potentially indicating more accelerated disease progression in DS.

### 2.3. DS individuals develop unique strains of Aβ with advanced AD, which differ in amounts of some tau and Aβ species

Environment-sensitive fluorescent dyes are ideal sensors for amyloid conformation because even small changes in local environment are exhibited in their emission spectra. While many such probes have been developed, we previously found a set of three dyes to sufficiently discriminate between AD-relevant Aβ strains in situ.^14^ We used this same set to examine the strains in this study, with optimized computational analysis (**Figure 3A**). Comparing these plaque-bound fluorescence spectra by PCA allowed us to identify structural differences between strains in high throughput (**Figure 3B–E**). Principle component 1 (PC1) represented 62% of the variation in the spectra, which was found to be due to a shelter-in-place-related microscope calibration change. We thus focused our analyses on PCs 2 and 3, which contained an additional 20% of the spectral variation. All three dyes contributed to the assessment (**Figure 3—figure supplement 1**). Using one-way MANOVA performed on patient centroid coordinates in PC2 and PC3, we determined that AD, DS, and ADNC are moderately but significantly differentiated by PCA (Wilks’ lambda [Λ]=0.74, p<0.005). AD-DS and DS in the absence of AD (here defined as *X*_tau_≤2) were similarly discriminated (Λ=0.69, p<0.005). This suggests that some the most prevalent Aβ strains in each stage of disease may be distinct.

**Figure 3.**
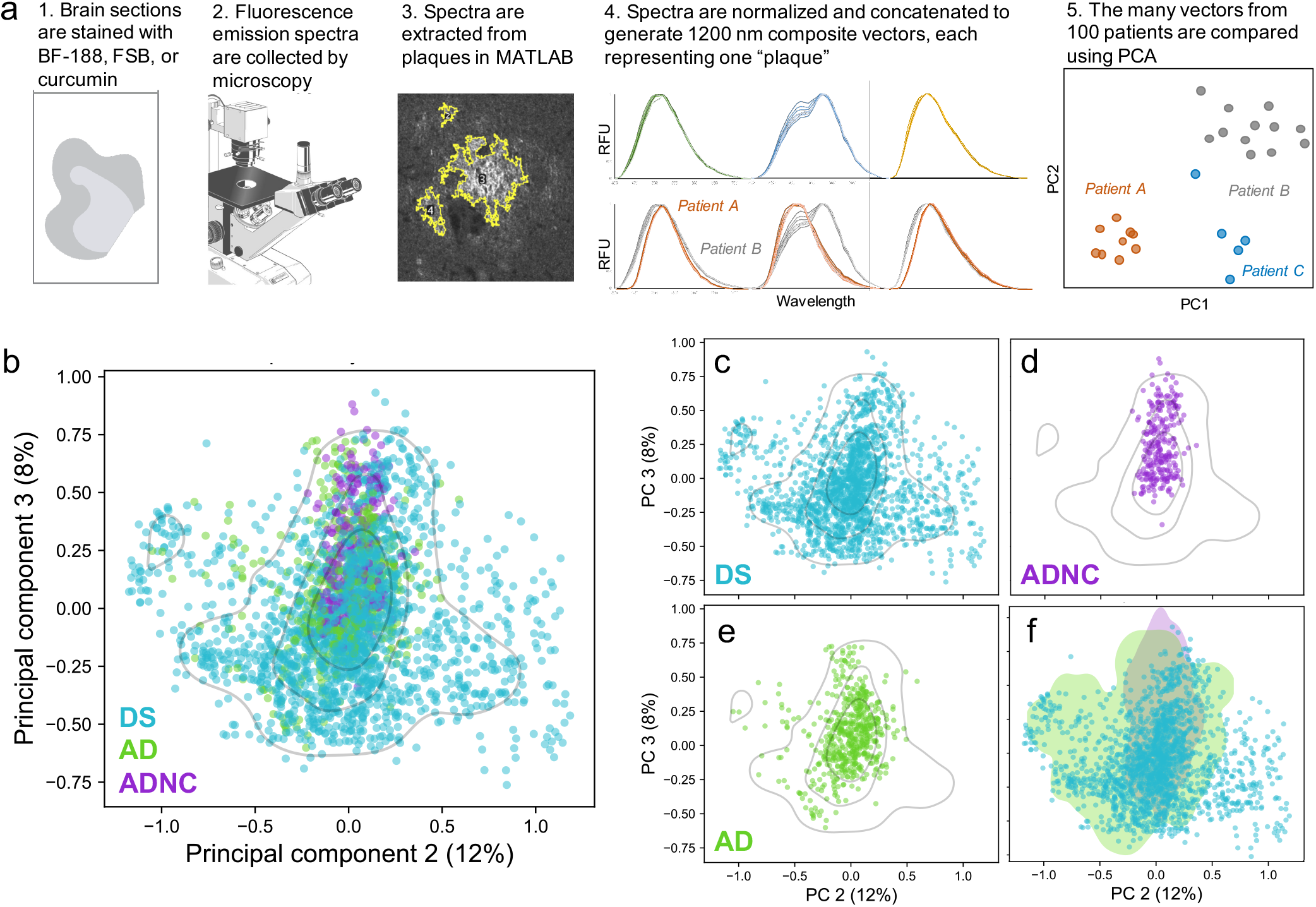
PCA performed on plaque-derived fluorescence spectra reveals a subset of conformational space unique to DS. (**A**) Overview of the experimental and computational workflow. (**B**) 2673 spectral vectors from 100 DS, AD, and ADNC cases were analyzed by PCA and are plotted in the eigenspace defined by PC2 and PC3. Each data point is the concatenated spectra of 3 dyes and represents one plaque. A Gaussian KDE is shown at 30%, 60%, and 90% probability intervals. (**C-E**) The PCA data are plotted separately by cohort for clarity. (**F)** A 99.5% KDE computed on either all ADNC vectors (purple) or all AD vectors (green) is shaded. The non-overlapping subset of DS vectors (blue points) indicate Aβ strains that may be unique to DS.

We defined two subsets of the eigenspace using a kernel density estimate (KDE) calculated on either all ADNC vectors or all AD vectors (**Figure 3F**). The overlap of the densities defines a region that contains plaques from all 3 cohorts. Considering that >95% of ADNC vectors are found in this region, these vectors could represent strains of Aβ that are found in normal, healthy aging, and could be less pathogenic. Interestingly, vectors from DS individuals aged 30-65 years, but not <30 years (n=3), are all found in this region, as are vectors from DS with both low (*X*_tau_=0-2) and high (*X*_tau_=4) AD pathology. The plaques not within these AD or ADNC densities indicate strains which, by the resolution of our method, are found exclusively in DS. These strains were found exclusively in individuals with high levels of pathology (*X*_tau_≥3).

In performing linear regression on patient centroids in each PC, we determined that the distribution of patients in PC2 is somewhat correlated with HTRF-measured [Aβ42] (r=0.34, p<0.05) in DS, whereas PC3 is significantly correlated with [Aβ40] (r=−0.36, p<0.005) when considering all patients. Alternatively, the concentrations of soluble, total, or pTau species did not significantly impact vector distributions in the eigenspace, nor did patient sex, age, or the tissue’s post-mortem interval (PMI) or source bank.

### 2.4. Strain heterogeneity increases with pathology in DS

We previously observed in sAD and fAD patients that the heterogeneity of Aβ strains varies both between populations and individuals.^14^ We were therefore curious how heterogeneous Aβ strains are in DS compared to sAD and ADNC individuals. To get an overall measure of the spread of patients in each cohort, we calculated variance-weighted RMSDs of the distances of each patient centroid to the cohort centroid. We found that DS patients were more heterogeneous (RMSD = 0.055) than AD patients (RMSD = 0.030), which were more heterogenous than ADNC patients (RMSD = 0.023) in this eigenspace, perhaps suggesting a greater difference in Aβ strains among DS individuals than among others.

Examples of per-patient vector populations are shown in **Figure 4A**. To quantify per-patient heterogeneity, we also calculated the RMSD of the distances of the patients’ vectors to their centroid. We found that like previously seen in fAD, per-patient RMSDs varied widely between patients but that strains were generally more homogeneous within a patient than for the entire population. In general, ADNC individuals were more homogeneous than AD cases, which were more homogenous than DS cases (**Figure 4B**). The proportion of DS cases with high RMSD were substantially greater in cases with advanced disease (*X*_tau_=3 or 4; **Figure 4C**) and age (**Figure 4—figure supplement 1A**). This suggests that the continued accumulation of Aβ in DS may result in its adoption of new or additional conformations. The presence of two *APOE* ε4 alleles, but not patient sex, also contributed to heterogeneity (**Figure 4—figure supplement 1B-C**).

**Figure 4.**
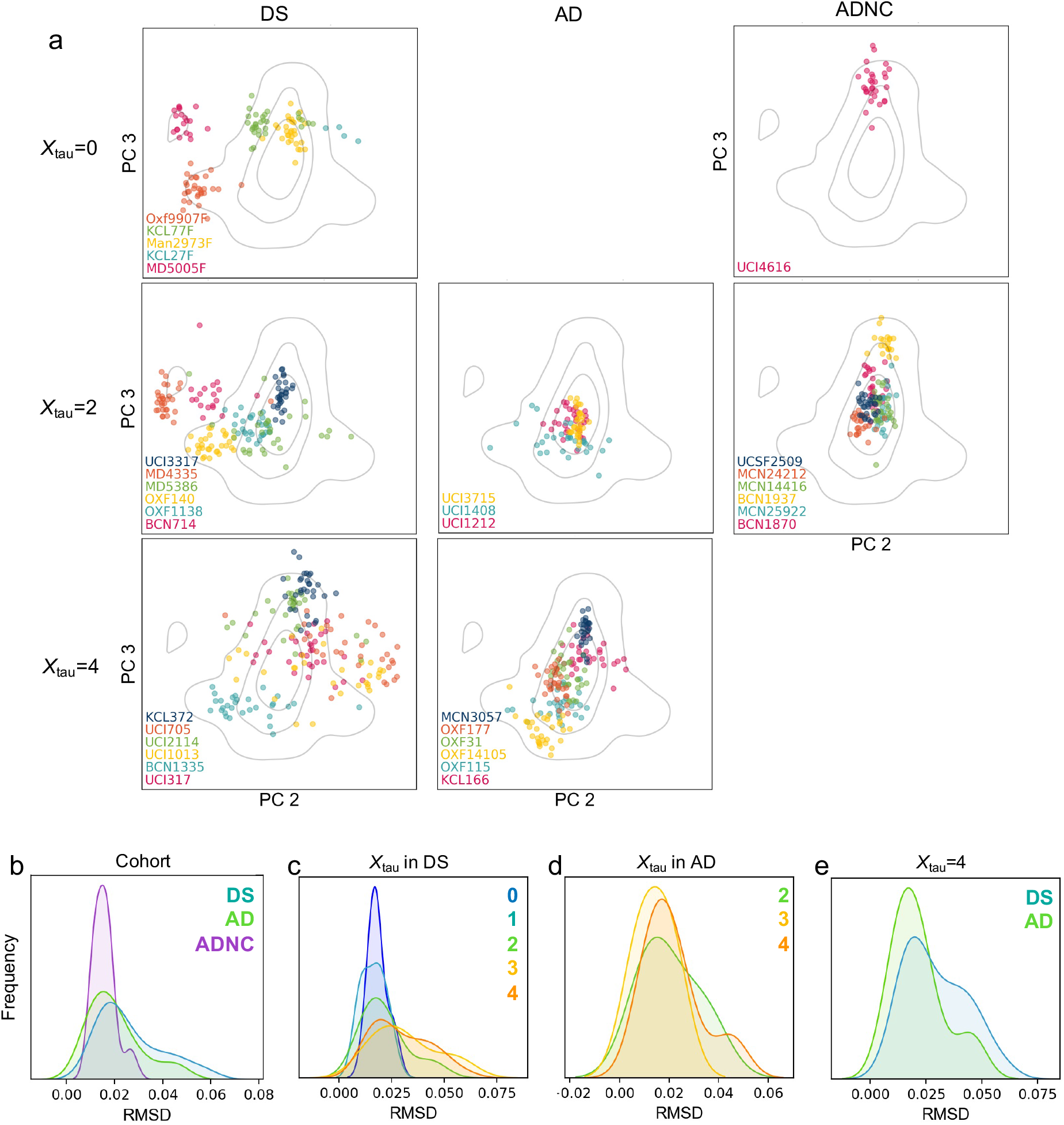
Per-patient strain heterogeneity increases with advancing pathology. (**A**) Examples of per-patient vector distributions in PC space. The columns show plots of DS, AD, and ADNC cases respectively. The rows show plots of cases with different *X*_tau_ in each cohort, when such cases exist. Each point is the spectral vector representing a single plaque from a given patient. (**B-E**) KDEs of RMSDs calculated using each case’s vector distributions in PCs 2-4. These distributions are compared between (**B**) cohorts, (**C**) *X*_tau_ in DS, (**D**) *X*_tau_ in AD, and (**E**) AD and DS cases with *X*_tau_=4.

## 3. DISCUSSION

### 3.1. Key neuropathological and biochemical differences distinguish AD-DS from sAD

Quantifying the progression of neuropathological hallmarks of sAD and AD-DS is critical for a comprehensive understanding of how Aβ and tau species might contribute—either independently or interactively—to the development of disease in each population. In DS, Aβ accumulation is known to begin in the late teenage years in the temporal lobe, with pathology developing throughout the brain in a similar pattern as in sAD by age 55.^28^ By IHC, we observed the earliest signs of Aβ deposition in DS as diffuse Aβ42 in the frontal cortices of individuals just younger than 10 years of age. In agreement with established trends,^46–48^ Aβ40 and Aβ42 both generally increased with age across the DS population, though Aβ40 was always found in the presence of appreciable Aβ42 accumulation. However, by HTRF, only Aβ40 and not Aβ42 levels were significantly elevated in DS over 40 years of age.

We also found slightly elevated levels of S202/T205 pTau by HTRF in AD-DS compared to AD and ADNC, which may indicate accelerated or more severe disease in DS. It is also interesting to note that while HTRF did not unveil significant differences in this species of pTau between AD and ADNC individuals, our neuropathological scoring method, which used an antibody against S262 pTau, greatly differentiated the two cohorts. A number of differences in the preparations of the two measured materials could be responsible. However, it is also possible that the difference is a consequence of different profiles of pTau that associate differently with NFTs or disease severity in AD. Current efforts in dissecting the timing and composition of pathogenic tau^49^ should certainly be expanded to include AD-DS to better understand the relationship between tau and disease.

Our finding the presence of Aβ in the frontal cortex as early as 9 years and some tau pathology as young as 19 years old in DS is earlier some others have reported.^25,28,47,50^ Since tau pathology does not extend beyond the temporal cortex to other regions of the cerebral cortex until Braak stage IV,^51^ it is also surprising that we found tau pathology in the frontal cortex of many cases assigned Braak stages I–III. These findings likely point to the sensitivity advantage of fluorescence-based methods over chromogenic stains. Yet overall, our neuropathological analysis highlights the heterogeneity of the DS brain, particularly in the relationships between Aβ and tau pathology, which do not appear to perfectly mirror that in sAD. This suggests that while on the whole AD-DS is an apt model for sAD, caution should be taken in assuming that all DS individuals are equally appropriate.

### 3.2. One subset of early-stage strains in DS reflect those in AD, but another subset is distinct

Considering the failure of multiple clinical trials for AD targeting the production of Aβ,^52^ it is critical that we recognize the diversity of Aβ species present in the brain and understand which, if any, might be most responsible for the onset or severity of dementia. To this end, we were interested in evaluating whether a unique subset of Aβ strains is present at early stages of AD, and whether early-stage strains persist or evolve with time or worsening clinical condition. To assess such relationships among strains in post-mortem tissue, we used environment-sensitive dyes to probe the amyloid plaques of individuals with DS and/or AD, as well as cognitively healthy ADNC individuals. We discriminated the plaque-bound fluorescence spectra using PCA, which successfully resolved strains of Aβ that are present at different stages of AD progression in DS. While these probes might report on other aspects of the plaque composition or environment,^53^ their specificity for insoluble protein aggregates and sensitivity to Aβ structure has been well documented^14,39,54,55^ and is expected to represent the majority of spectral variation.

We found that some of the strains we observed in the frontal cortices of DS individuals with minimal to non-existent NFTs but some Aβ pathology, i.e. generally before the onset of clinical symptoms of AD, are distinct from strains in AD-DS. In DS cases where tau pathology was more severe, many strains appeared common to AD and AD-DS. Intriguingly, a portion of plaques in many late-stage AD-DS individuals appear distinct from any non-DS plaques, suggesting that not all strains of Aβ are common to the two diseases. However, these cross-sectional observations can only suggest what may be happening temporally in these individuals. Moving forward, using strain-specific imaging agents in live subjects with DS will be critical to understanding the true evolution of Aβ strains in AD.

While the difference in amyloid presentation in senile plaques itself may be important for the development of anti-amyloid AD therapeutics, the causes of these different strains may also inform our understand of AD in DS. For example, neuroinflammation, in particular plaque-associated microglia activation, is known to affect amyloid conformation.^56–58^ As the relative populations of different types of microglial cells are altered in people with DS after the age of 40,^59,60^ these changes may manifest in altered amyloid conformation. Perhaps most critically, the progressive changes in strain composition evinced by PCA may indicate that distinct strains of Aβ are present prior to the onset of clinical AD. It is possible that these strains persist while other strains emerge along with clinical symptoms, in which case the next obvious question arises: are these strains harmless bystanders, or are they the catalyst of disease? If the latter, it would be critical to ensure that preventative therapies and diagnostics targeted these strains.

### 3.3. Aβ strain heterogeneity increases with AD progression

Through calculating the RMSDs of case vector distributions in principal component space, we have found that Aβ strain heterogeneity varies among individuals. Heterogeneity tended be greater in individuals with more advanced AD, particularly in DS. In most of these DS individuals, we found representation of strains both common to and unique from late-stage sAD. This suggests that while many strains common to sAD and AD-DS persist throughout life, certain conditions specific to aging or pathology in DS may allow for the emergence of new strains. Recent evidence shows that while APP has a direct role in AD-related Aβ neuropathology,^61,62^ other genes on Chr21 also substantially impact Aβ aggregation in AD-DS^63^. As tau is thought to interact with Aβ in AD,^4^ changes in tau isoform ratios and phosphorylation via Chr21-associated proteins DYRK1A^64,65^ and RCAN1^66^ could also impact Aβ plaque composition. Our finding of enhanced pTau in DS individuals over 40 years of age supports this hypothesis. These DS-specific changes could therefore enhance this strain diversification or alter the specific strains that are favored.

The possibility that multiple distinct Aβ species or a spectrum of species could exist within a single patient has important therapeutic implications. For example, a changed cellular environment could favor the dominant propagation of a previously minor strain. Should this strain be particularly pathogenic and negatively impact other neuropathological or clinical outcomes, then preventing the emergence of such a strain would be of paramount therapeutic importance. In the worst-case scenario, a seemingly successful treatment aimed nonspecifically at Aβ could be thwarted by the evolution of drug resistant strains, as has been observed in screens against prions causing scrapie.^67,68^ Furthermore, targeting only one or a few of many pathogenic strains in a given individual could have little impact on slowing disease progression—an outcome possibly demonstrated in the failure of anti-amyloid therapies.^52,69^

Understanding the structural characteristics of these strains in more detail will help us understand how they may evolve and what role they may play in AD in DS. Ultimately, we might not be able to treat DS as a direct model for all sAD. Instead, by understanding what features in DS might be associated with altered Aβ strain profiles, we could triage clinical trial candidates accordingly. Based on the findings presented here, we would suggest that DS individuals already have Aβ conformations or plaque environments that replicate those in AD before the onset of dementia, which can differentiate with age and advancing disease. Thus, we posit that younger, non-demented individuals with DS may be the only appropriate DS candidates for clinical trials targeting sAD, while more pathologically advanced individuals would require a separate therapeutic strategy.

### 3.4. Future work

Whether the unique strains developing late in AD-DS are emergent or native to the individual is still not clear. A next important step will be to assess how these strains might differ among brain regions, particularly in the hippocampus where Aβ is believed to spread from the neocortex.^70,71^ Machine learning could also be applied to assess morphological and intra-plaque differences in our existing micrographs in order to more robustly differentiate strains. The development of PET imaging agents that are specific to multiple distinct Aβ strains will certainly be needed in order to follow these potential changes longitudinally. Furthermore, the success of future AD diagnostic and therapeutic efforts against Aβ depends on a detailed structural understanding of these strains. Mass spectrometry and cryo-EM or ssNMR should be employed to precisely understand the commonalities between specific strains in sAD and AD-DS, and moreover what aspects of plaque composition or amyloid structure make certain strains unique.

### 3.5. Conclusion

This work provides the first analysis of Aβ strains in DS and their relevance to sAD. We showed that AD-DS generally reflects the broad neuropathological features of AD but differs significantly in Aβ40 and pTau concentrations. Through molecular analysis using environment-sensitive fluorescent probes, we found that DS, AD-DS, AD, and ADNC all likely share a subset of Aβ strains. However, as AD progresses in DS, strains become more heterogeneous and some prominent strains tend to diverge from non-AD-like Aβ. We therefore suggest that AD clinical trials focus on recruitment of younger DS patients who do not yet show signs of dementia; in doing so, however, it must be recognized that more heterogenous dominant strains of Aβ in AD-DS would potentially not be yet recognized. It is critical to follow this work with high-resolution structural analysis of the differences between Aβ strains in older and younger DS and to understand the mechanistic connections between the DS brain environment and Aβ heterogeneity.

## 4. METHODS

### 4.1. Cases

Deidentified post-mortem brain tissue was obtained from 210 individuals: 152 DS (+/- AD), 34 AD without DS, and 24 control cases without cognitive impairment but with AD neuropathological change (ADNC). Details on each case are outlined in **Table 1** (attached). Included in the DS cohort is one subject with partial trisomy of Chr21 (PT21) that does not include *APP*, which resulted in normal aging without dementia^62^ and affords us interesting comparisons between characteristics that might differ in DS without the eventuality of AD. Frozen blocks and/or formalin-fixed paraffin-embedded (FFPE) sections were analyzed from the frontal cortex. Note that not all cases were able to be used for every experiment, depending on amount and type of tissue preparation available for each case (see **Table 1**). All cases able to be used in an experiment are included in the presented results unless otherwise specified.

### 4.2. Immunohistochemistry (IHC)

Deparaffinized fixed sections were pretreated in 98% formic acid for 6 min to enhance immunoreactivity. After blocking with 10% normal goat serum (ngs) in PBS with 0.2% Tween 20 (PBST), sections were incubated at room temperature in primary antibodies overnight followed by secondary antibodies for 2 h. Primaries were prepared in 10% ngs and applied as combinations of either: anti-Aβ1-40 rabbit polyclonal (Millipore Sigma #AB5074P, 1:200) and anti-Aβ1-42 12F4 mouse monoclonal (Biolegend #805502, 1:200); or anti-Aβ17-24 4G8 mouse monoclonal (Biolegend #800709 1:1000) and anti-tau (phospho-S262) rabbit polyclonal (Abcam #ab131354, 1:200). Polyclonal IgG H&L secondaries were Alexa Fluor 488- and 647-conjugates (Thermo Fisher #s A11029, A21235, A11008, and A21244) applied 1:500 in 10% ngs in PBST.

Stained slides were scanned on a ZEISS Axio Scan Z1 digital slide scanner at 20x magnification. Excitation at 493, 553, and 653 nm was followed by detection at 517 nm (Aβ40 or phospho-tau), 568 nm (autofluorescence), and 668 nm (Aβ42 or tau).

### 4.3. Neuropathological scoring

To determine the level of AD pathology at the time of death, one IHC-stained fixed cortical section was evaluated for each case. The number of Aβ40- and Aβ42-positive plaques, neuritic plaques, and phospho-S262-positive NFTs were averaged among three random 1-mm^2^ sections of grey matter. Aβ and tau scores were assigned according to the criteria in **Table 2**, which were formulated to honor traditional staging methods, to allow for scorer efficiency, and to separate the patient pool into large enough groups to facilitate downstream analysis.

NFT accumulation in the neocortex (Braak stage V-VI) is required for a post-mortem diagnosis of AD. Therefore, to cross-check any assignment of *X*_tau_=0 in AD, a BF-188-stained fixed section was viewed with a red-light filter by confocal microscopy, which would reveal both phosphorylated and unphosphorylated tau species^54^. We eliminated from downstream analysis AD cases that indeed appeared to have no NFTs (n=2) in the frontal cortex.

### 4.4. DNA extraction and genotyping

Upon receipt, frozen brain tissue was homogenized in PBS and stored 10% w/v in PBS at −80 °C until thawed on ice for biochemical assays. Genomic DNA was purified from this homogenate using a DNeasy Blood & Tissue Kit (Qiagen cat #69506).

To determine the *APOE* genotype of each case, gene fragments encompassing the two APOE-relevant SNPs were amplified by PCR based on the protocol described by Zhong et al. 2016.^72^ Each 50-μL reaction contained 1 U Phusion Plus DNA Polymerase, 200 μM dNTPs, 1X Phusion GC buffer, 5% DMSO, 0.2 μM forward and reverse primers, and 10-100 ng gDNA. Primer sequences and cycling conditions are in **Supplementary Table 1**. Purified PCR products were Sanger sequenced by Genewiz (San Francisco, CA).

### 4.5. Protein quantification

To determine the total concentration of soluble APP and tau present in each frontal cortex sample, sandwich enzyme-linked immunosorbent assays (ELISAs; Invitrogen, cat #s KHB0051 and KHB0041) were performed on brain homogenate (10% in PBS, called “10% BH”) clarified through centrifugation to remove cell debris and the majority of insoluble proteins. Samples were prepared and stored in low-binding 96-well plates and measured according to manufacturer directions. It should be noted that a subset of samples from two tissue banks were measured separately in time and were found to have 10-100x less soluble tau than the lowest other sample; this set of samples was not included in any bulk analyses on the assumption of batch error. Protein concentrations were normalized to total brain protein in the clarified homogenate as determined by BCA.

Insoluble protein fractions were extracted from brain homogenate by sonicating 10% BH with 75% v/v formic acid for 20 min followed by ultracentrifugation at 48000*x*g for 1 h at 4 °C. The supernatant was neutralized with 20-fold dilution in neutralization buffer (1M tris base [NH_2_C(CH_2_OH)_3_] 0.5M Na_2_HPO_4_ · 7H_2_O, pH 10.5) and was stored in aliquots at −80 °C until use. To measure concentrations of Aβ40, Aβ42, and insoluble tau species in these extracts, ELISAs were attempted but were abandoned due to the imprecision of biological replicates. Therefore, homogeneous time-resolved fluorescence (HTRF) assays were performed instead. Total tau (Perkin Elmer Cisbio 64NTAUPEG), tau phospho-S202/T205 (64TS2PEG), Aβ40 (62B40PEG), and Aβ42 (62B42PEG) HTRF kits were used according to manufacturer protocols. Peptide standards were not provided in either tau kit for generating standard concentration curves, so unphosphorylated and hyperphosphorylated 0N4R tau from insect cell expressions (gifts from Aye Thwin and Dr. Greg Merz, UCSF) were used after optimization of standard concentration ranges.

### 4.6. Spectral profiling of plaque-bound fluorescent dyes

Cases with *X*_Aβ_=0, including the PT21 case, could not undergo Aβ strain analysis due to their lack of plaques. For cases with sufficient pathology, adjacent cortical sections were blocked in PBST, stained with 2.5 μM curcumin, BF-188, and FSB prepared in PBS (with 5% EtOH for curcumin), and washed in PBS. Labelled plaques were imaged in the spectral (Lambda) scan mode of a Leica SP8 confocal microscope using a 40x water immersion lens (1.1 NA), a 405-nm laser for excitation, and a HyD detector at 512- × 512-pixel resolution. For each field-of-view, the optical plane was moved to the center of the z-stack volume for a given Aβ deposit, and fluorescence emission was acquired from a series of 40-image steps spanning 385- to 780-nm wavelengths using a sliding 15-nm-wide detection window.

Micrographs were analyzed using custom MATLAB^73^ software. Plaques were automatically segmented based on size and fluorescence intensity. False-positive objects, including neuritic plaque-associated NFTs, were manually excluded to ensure that fluorescence information was plaque-specific. Separate spectra obtained from each of the three dyes were normalized to their maximum intensities and randomly concatenated to form the full 1200-nm spectral vector for each patient case used in PCA. To avoid biasing PCA towards individuals with more plaques, we limited the analysis to 30 randomly-selected vectors per patient. We compared multiple under-sampled PCA to each other and to PCA that included all possible vectors to ensure that the trends observed in each were the same.

### 4.7. PCA and statistical analysis

PCA was performed on composite spectral vectors using the Python sklearn decomposition package. Comparisons made between groups were always performed in the same eigenspace. Density contours were applied to the PCA plot using the matplotlib contour function calculated on a Gaussian KDE mesh grid within the Scipy stats package. To account for the overrepresentation of DS cases in our analysis, we validated our PCA through computational oversampling of young DS cases as well as through the inclusion of all (>30) spectra from the given AD cases, neither of which altered the relationships among samples in PC space.

To determine the heterogeneity of strains, the weighted root-mean-square deviation (RMSD) of spectral vectors in principle components (PCs) 2, 3, and 4 was calculated for each patient and for each cohort using the following equation:

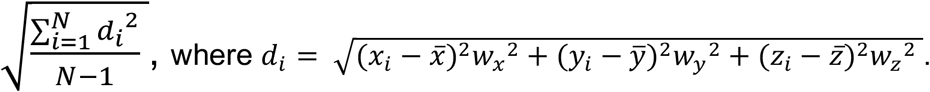

Per-patient calculations were performed on each patient vector with *N* as the total number of patient spectral vectors and the barred coordinates representing the patient centroid. Per-cohort calculations were calculated on the centroid of each patient in the cohort with *N* as the total number of patients in the cohort and the barred coordinates representing the cohort centroid. The weights are the proportion of overall variance explained by PCs 2, 3 and 4.

Linear regression from SciPy and the StatAnnot package were used to determine significance of relationships between PCA coordinates and numerical and categorical case attributes respectively. One-way multivariate analysis of variance (MANOVA) was performed on the centroids of each patient in PC2 and PC3 to determine the resolving power of PCA among groups.

## 5. ACKNOWLEDGEMENTS

We are grateful to Drs. Elizabeth Head (UCI) and William Mobley (UCSD) for their thoughtful advice; to John Nicoludis (Invitae), Greg Merz (UCSF), Robert Newberry (UCSF), and Bruk Mensa (UCSF) for their technical guidance; and to Abby Oehler and Bailin Ye for their technical assistance. This material is based on work supported by the National Science Foundation Graduate Research Fellowship Program under Grant No. 2034836 and by the NIH under Grants Nos. T32 GM064337 (A.M.M), RF1AG061874 (C.C. and W.F.D.) and P01AG002132 (C.C. and W.F.D.). Any opinions, findings, and conclusions or recommendations expressed in this material are those of the author(s) and do not necessarily reflect the views of the NSF or NIH. Tissue samples were supplied by the NIH NeuroBioBank; the August Pi i Sunyer Biomedical Research Institute (IDIBAPS) Biobank (Barcelona, Spain); the University of California Alzheimer’s Disease Research Center (UCI-ADRC, Irvine, CA), which is funded by NIH/NIA Grant P30AG066519; the UCSF Neurodegenerative Disease Brain Bank and Prof. William W. Seeley (UCSF Memory and Aging Center, San Francisco, CA); the University of Washington Neuropathology Core (Seattle, WA), which is supported by the Alzheimer’s Disease Research Center (AG005136), the Adult Changes in Thought Study (AG006781), and the Morris K. Udall Center of Excellence for Parkinson’s Disease Research (NS062684); the London Neurodegenerative Diseases Brain Bank (King’s College London, England), which receives funding from the Medical Research Council UK and through the Brains for Dementia Research Project (jointly funded by the Alzheimer’s Society and Alzheimer’s Research UK); the Oxford Brain Bank, supported by the Medical Research Council (MRC), Brains for Dementia Research (BDR) (Alzheimer Society and Alzheimer Research UK), Autistica UK and the NIHR Oxford Biomedical Research Centre; the Langone Health Alzheimer’s Disease Center (New York University, NY), which is supported by funding from NIH grant to the NYU Alzheimer’s Disease Research Center, P30AG066512; and The Manchester Brain Bank (University of Manchester, England), which is part of the Brains for Dementia Research program, jointly funded by Alzheimer’s Research UK and Alzheimer’s Society.

## 6. COMPETING INTERESTS

The authors declare no competing interests.

## 7. SUPPLEMENTARY FIGURES

**Figure 2—figure supplement 1:**
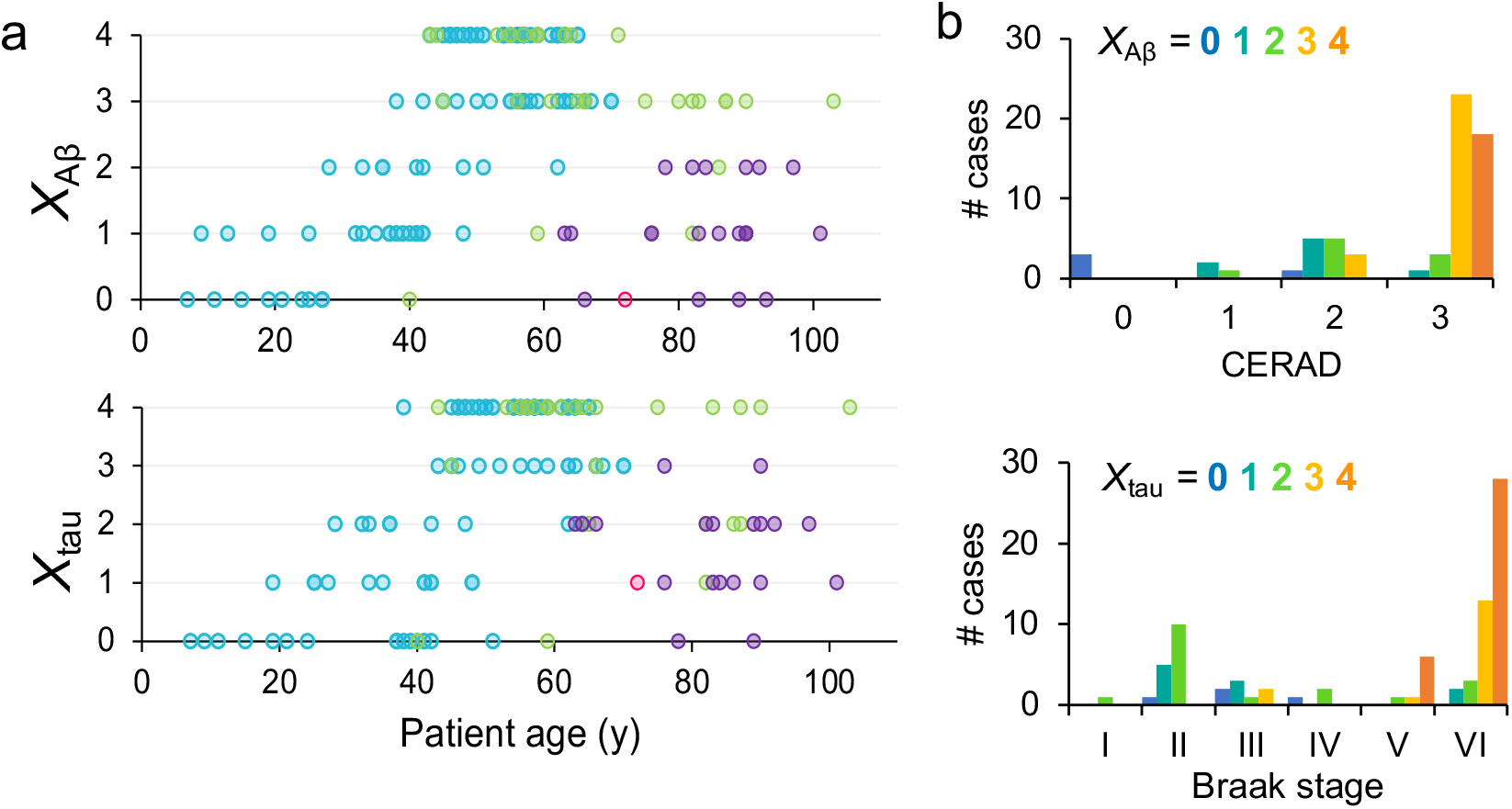
Neuropathology scores compared to age and autopsy scores. **(A)** Aβ and tau scores relative to patient age at death. DS subjects are shown in blue, AD in green, ADNC in purple, and PT21 in pink. (**B**) Relationship between manual Aβ and tau scores and bank-provided scores determined at autopsy.

**Figure 4—figure supplement 1:**
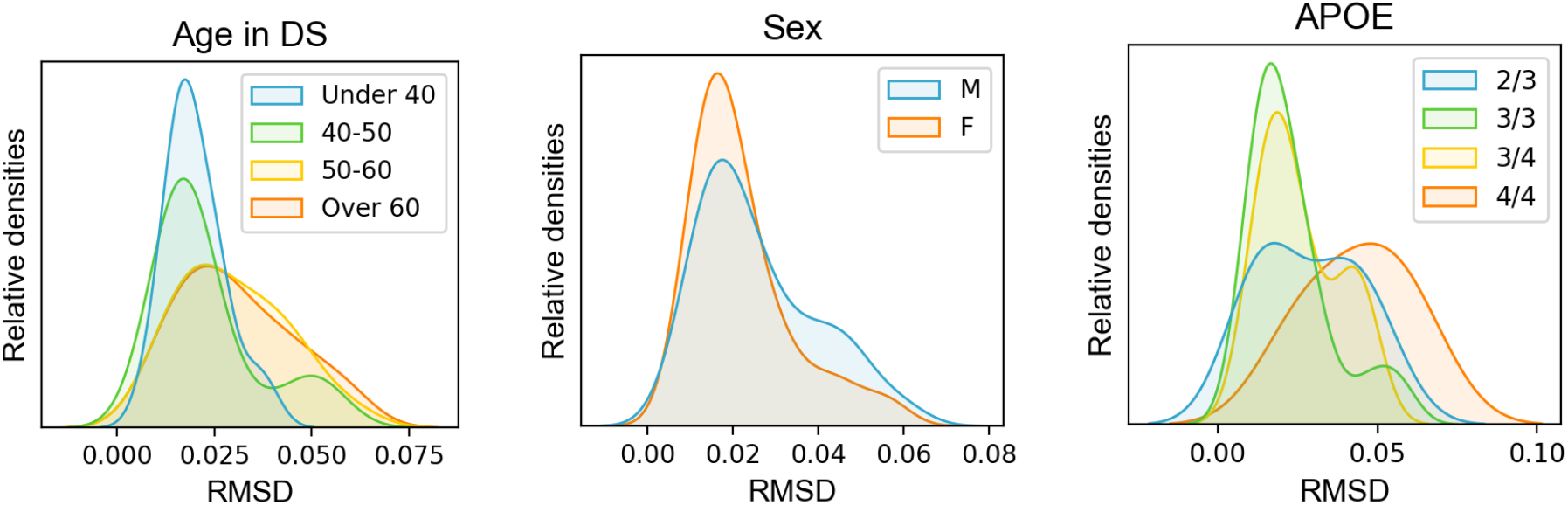
Per-patient RMSD distributions by age, sex, and APOE. (**A**) RMSD by DS patient age at death, (**B**) by sex, and (**C**) by APOE genotype, when known. RMSD values were calculated as the distances in PC2, PC3, and PC4 of each vector from the centroid of all the vectors for a given case. The relative densities are shown as a Gaussian KDE, normalized to the area under the curve, of the RMSDs calculated for all the cases in a given group.

**Figure 2—figure supplement 2:**
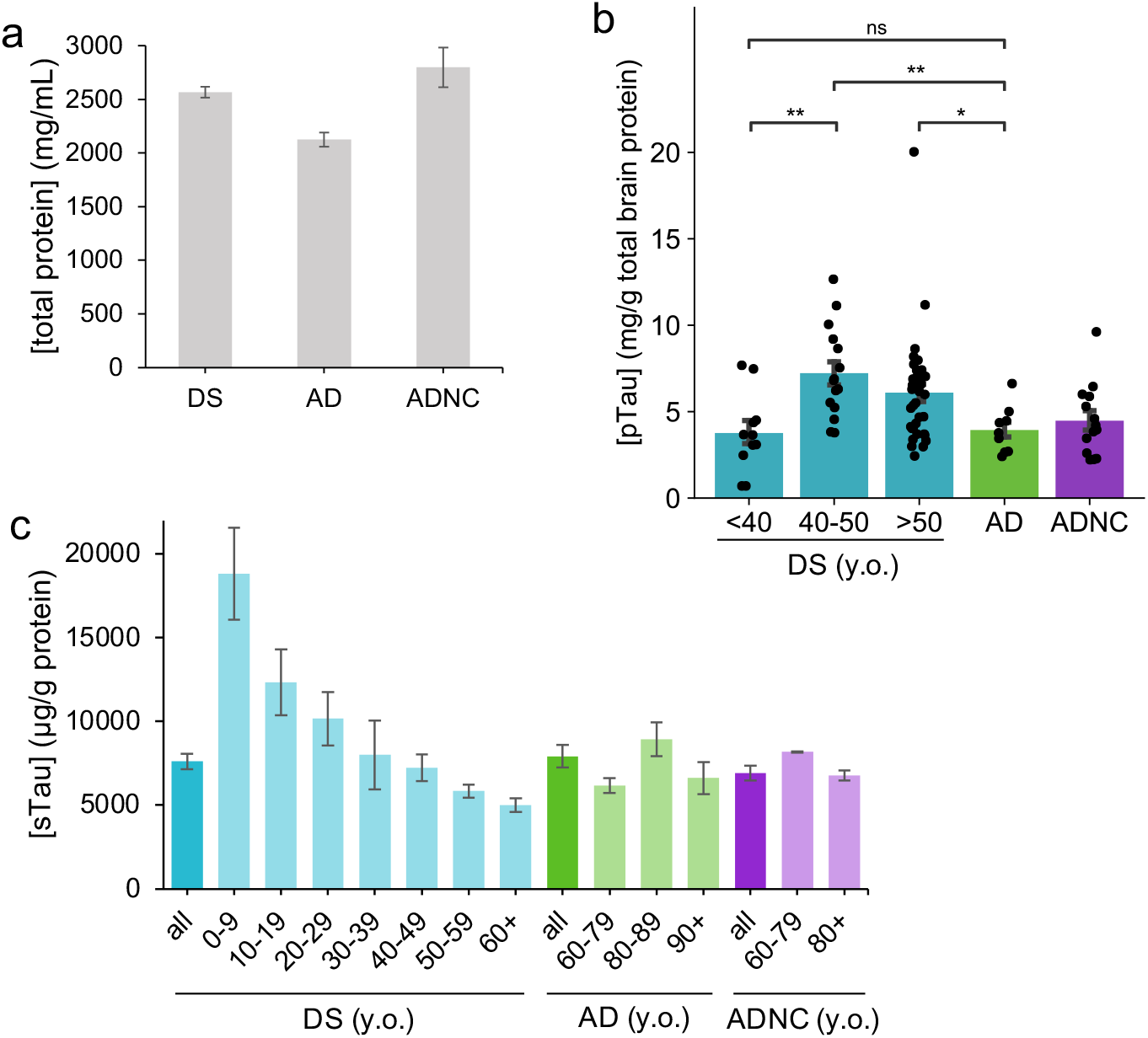
Total protein concentrations per cohort and tau species concentrations in greater granularity. (**A**) Total protein in 10% BH assayed by BCA (**B**) S203/T205 pTau concentrations as measured by HTRF delineated by age in DS, ± SEM. pTau is significantly higher in DS after 40 years of age compared to in younger DS individuals and compared to AD. *p*-value: *: 0.01 < p ≤ 0.05; **: 0.001 < p ≤ 0.01, ***: 0.0001 < p ≤ 0.001 (**C**) Soluble tau concentrations as measured by ELISA decline with age in DS, ± SEM.

**Figure 3—figure supplement 1:**
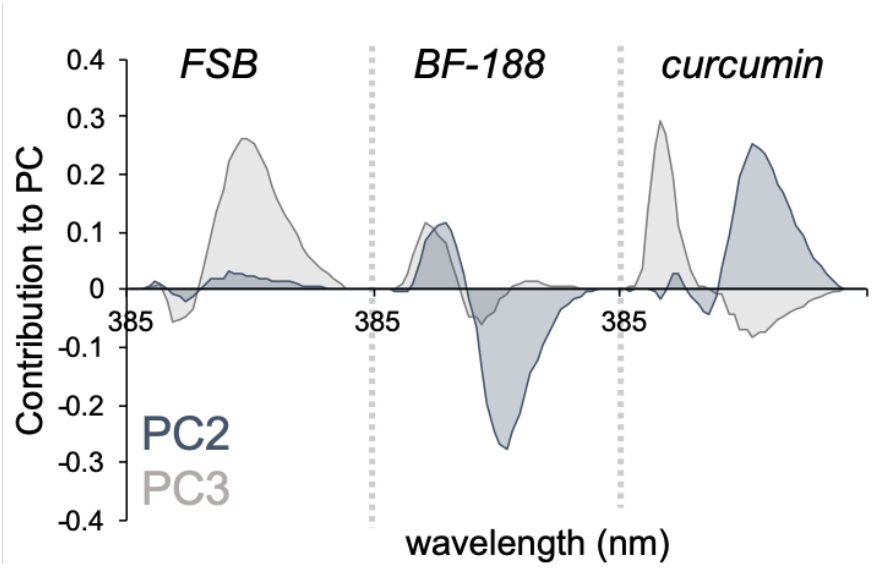
Wavelength contributions to the principle component space. Each of the three conformation-sensitive dyes FSB, BF-188, and curcumin contribute to PC2 and PC3.

## 8. SUPPLEMENTARY TABLES

**Supplementary Table 1.**
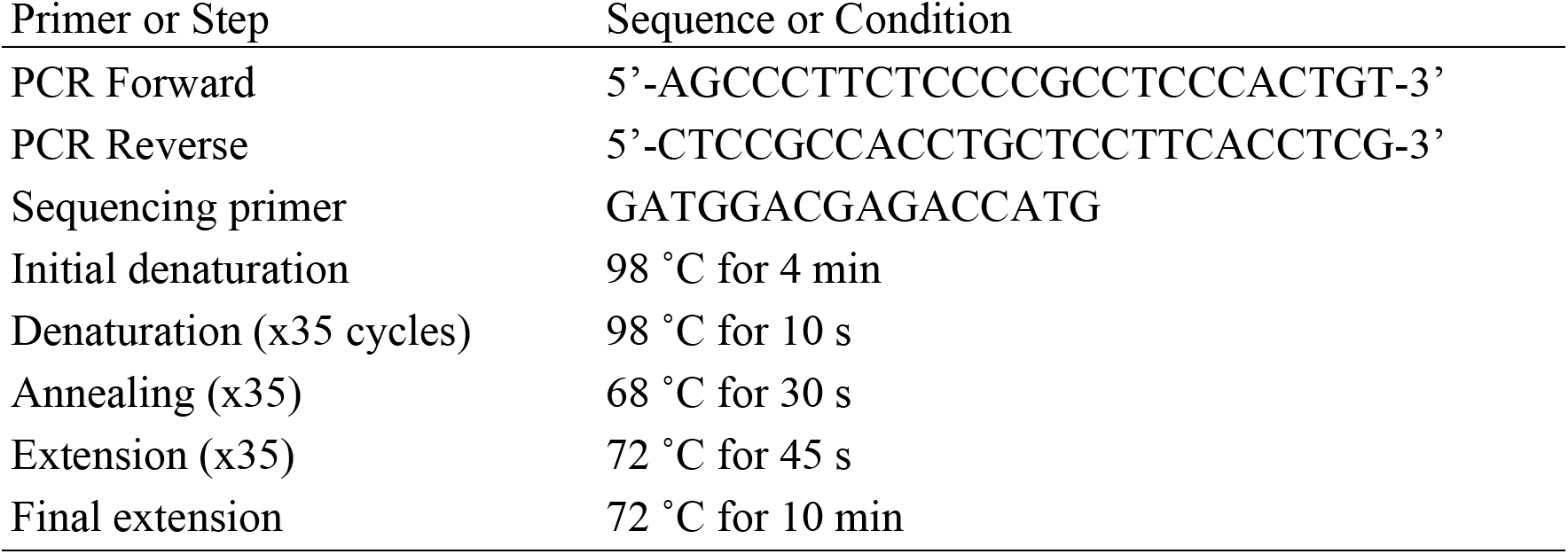
Primers and conditions used in PCR and sequencing.

## Notes

### Competing Interest Statement

The authors have declared no competing interest.

## REFERENCES

(1) Hebert, L. E.; Weuve, J.; Scherr, P. A.; Evans, D. A. Alzheimer Disease in the United States (2010–2050) Estimated Using the 2010 Census. Neurology 2013, 80 (19), 1778–1783. https://doi.org/10.1212/WNL.0b013e31828726f5.

(2) Kumar, A.; Singh, A.; Ekavali. A Review on Alzheimer’s Disease Pathophysiology and Its Management: An Update. Pharmacol. Rep. 2015, 67 (2), 195–203. https://doi.org/10.1016/j.pharep.2014.09.004.

(3) 2020 Alzheimer’s Disease Facts and Figures. Alzheimers Dement. 2020, 16 (3), 391–460. https://doi.org/10.1002/alz.12068.

(4) Nisbet, R. M.; Polanco, J.-C.; Ittner, L. M.; Götz, J. Tau Aggregation and Its Interplay with Amyloid-β. Acta Neuropathol. (Berl.) 2015, 129, 207–220. https://doi.org/10.1007/s00401-014-1371-2.

(5) Thal, D. R.; Rüb, U.; Orantes, M.; Braak, H. Phases of A Beta-Deposition in the Human Brain and Its Relevance for the Development of AD. Neurology 2002, 58 (12), 1791–1800. https://doi.org/10.1212/wnl.58.12.1791.

(6) Villemagne, V. L.; Burnham, S.; Bourgeat, P.; Brown, B.; Ellis, K. A.; Salvado, O.; Szoeke, C.; Macaulay, S. L.; Martins, R.; Maruff, P.; Ames, D.; Rowe, C. C.; Masters, C. L.; Australian Imaging Biomarkers and Lifestyle (AIBL) Research Group. Amyloid β Deposition, Neurodegeneration, and Cognitive Decline in Sporadic Alzheimer’s Disease: A Prospective Cohort Study. Lancet Neurol. 2013, 12 (4), 357–367. https://doi.org/10.1016/S1474-4422(13)70044-9.

(7) Bierer, L. M.; Hof, P. R.; Purohit, D. P.; Carlin, L.; Schmeidler, J.; Davis, K. L.; Perl, D. P. Neocortical Neurofibrillary Tangles Correlate With Dementia Severity in Alzheimer’s Disease. Arch. Neurol. 1995, 52 (1), 81–88. https://doi.org/10.1001/archneur.1995.00540250089017.

(8) Bejanin, A.; Schonhaut, D. R.; La Joie, R.; Kramer, J. H.; Baker, S. L.; Sosa, N.; Ayakta, N.; Cantwell, A.; Janabi, M.; Lauriola, M.; O’Neil, J. P.; Gorno-Tempini, M. L.; Miller, Z. A.; Rosen, H. J.; Miller, B. L.; Jagust, W. J.; Rabinovici, G. D. Tau Pathology and Neurodegeneration Contribute to Cognitive Impairment in Alzheimer’s Disease. Brain 2017, 140 (12), 3286–3300. https://doi.org/10.1093/brain/awx243.

(9) Joie, R. L.; Visani, A. V.; Baker, S. L.; Brown, J. A.; Bourakova, V.; Cha, J.; Chaudhary, K.; Edwards, L.; Iaccarino, L.; Janabi, M.; Lesman-Segev, O. H.; Miller, Z. A.; Perry, D. C.; O’Neil, J. P.; Pham, J.; Rojas, J. C.; Rosen, H. J.; Seeley, W. W.; Tsai, R. M.; Miller, B. L.; Jagust, W. J.; Rabinovici, G. D. Prospective Longitudinal Atrophy in Alzheimer’s Disease Correlates with the Intensity and Topography of Baseline Tau-PET. Sci. Transl. Med. 2020, 12 (524). https://doi.org/10.1126/scitranslmed.aau5732.

(10) Tanzi, R. E.; Bertram, L. Twenty Years of the Alzheimer’s Disease Amyloid Hypothesis: A Genetic Perspective. Cell 2005, 120 (4), 545–555. https://doi.org/10.1016/j.cell.2005.02.008.

(11) Goedert, M.; Crowther, R. A.; Spillantini, M. G. Tau Mutations Cause Frontotemporal Dementias. Neuron 1998, 21 (5), 955–958. https://doi.org/10.1016/S0896-6273(00)80615-7.

(12) Lam, B.; Masellis, M.; Freedman, M.; Stuss, D. T.; Black, S. E. Clinical, Imaging, and Pathological Heterogeneity of the Alzheimer’s Disease Syndrome. Alzheimers Res. Ther. 2013, 5 (1), 1. https://doi.org/10.1186/alzrt155.

(13) Rasmussen, J.; Mahler, J.; Beschorner, N.; Kaeser, S. A.; Häsler, L. M.; Baumann, F.; Nyström, S.; Portelius, E.; Blennow, K.; Lashley, T.; Fox, N. C.; Sepulveda-Falla, D.; Glatzel, M.; Oblak, A. L.; Ghetti, B.; Nilsson, K. P. R.; Hammarström, P.; Staufenbiel, M.; Walker, L. C.; Jucker, M. Amyloid Polymorphisms Constitute Distinct Clouds of Conformational Variants in Different Etiological Subtypes of Alzheimer’s Disease. Proc. Natl. Acad. Sci. 2017, 114 (49), 13018–13023. https://doi.org/10.1073/pnas.1713215114.

(14) Condello, C.; Lemmin, T.; Stöhr, J.; Nick, M.; Wu, Y.; Maxwell, A. M.; Watts, J. C.; Caro, C. D.; Oehler, A.; Keene, C. D.; Bird, T. D.; van Duinen, S. G.; Lannfelt, L.; Ingelsson, M.; Graff, C.; Giles, K.; DeGrado, W. F.; Prusiner, S. B. Structural Heterogeneity and Intersubject Variability of Aβ in Familial and Sporadic Alzheimer’s Disease. Proc. Natl. Acad. Sci. 2018, 201714966. https://doi.org/10.1073/pnas.1714966115.

(15) Condello, C.; Stöehr, J. Aβ Propagation and Strains: Implications for the Phenotypic Diversity in Alzheimer’s Disease. Neurobiol. Dis. 2018, 109 (Pt B), 191–200. https://doi.org/10.1016/j.nbd.2017.03.014.

(16) Colletier, J.-P.; Laganowsky, A.; Landau, M.; Zhao, M.; Soriaga, A. B.; Goldschmidt, L.; Flot, D.; Cascio, D.; Sawaya, M. R.; Eisenberg, D. Molecular Basis for Amyloid-β Polymorphism. Proc. Natl. Acad. Sci. 2011, 108 (41), 16938–16943. https://doi.org/10.1073/pnas.1112600108.

(17) Toyama, B. H.; Weissman, J. S. Amyloid Structure: Conformational Diversity and Consequences. Annu. Rev. Biochem. 2011, 80. https://doi.org/10.1146/annurev-biochem-090908-120656.

(18) Cohen, M. L.; Kim, C.; Haldiman, T.; ElHag, M.; Mehndiratta, P.; Pichet, T.; Lissemore, F.; Shea, M.; Cohen, Y.; Chen, W.; Blevins, J.; Appleby, B. S.; Surewicz, K.; Surewicz, W. K.; Sajatovic, M.; Tatsuoka, C.; Zhang, S.; Mayo, P.; Butkiewicz, M.; Haines, J. L.; Lerner, A. J.; Safar, J. G. Rapidly Progressive Alzheimer’s Disease Features Distinct Structures of Amyloid-β. Brain 2015, 138 (4), 1009–1022. https://doi.org/10.1093/brain/awv006.

(19) Qiang, W.; Yau, W.-M.; Lu, J.-X.; Collinge, J.; Tycko, R. Structural Variation in Amyloid-β Fibrils from Alzheimer’s Disease Clinical Subtypes. Nature 2017, 541 (7636), 217–221. https://doi.org/10.1038/nature20814.

(20) Watts, J. C.; Condello, C.; Stöhr, J.; Oehler, A.; Lee, J.; DeArmond, S. J.; Lannfelt, L.; Ingelsson, M.; Giles, K.; Prusiner, S. B. Serial Propagation of Distinct Strains of Aβ Prions from Alzheimer’s Disease Patients. Proc. Natl. Acad. Sci. 2014, 111 (28), 10323–10328. https://doi.org/10.1073/pnas.1408900111.

(21) Stohr, J.; Watts, J. C.; Mensinger, Z. L.; Oehler, A.; Grillo, S. K.; DeArmond, S. J.; Prusiner, S. B.; Giles, K. Purified and Synthetic Alzheimer’s Amyloid Beta (A) Prions. Proc. Natl. Acad. Sci. 2012, 109 (27), 11025–11030. https://doi.org/10.1073/pnas.1206555109.

(22) Stohr, J.; Condello, C.; Watts, J. C.; Bloch, L.; Oehler, A.; Nick, M.; DeArmond, S. J.; Giles, K.; DeGrado, W. F.; Prusiner, S. B. Distinct Synthetic AB Prion Strains Producing Different Amyloid Deposits in Bigenic Mice. Proc. Natl. Acad. Sci. 2014, 111 (28), 10329–10334. https://doi.org/10.1073/pnas.1408968111.

(23) Bateman, R. J.; Aisen, P. S.; De Strooper, B.; Fox, N. C.; Lemere, C. A.; Ringman, J. M.; Salloway, S.; Sperling, R. A.; Windisch, M.; Xiong, C. Autosomal-Dominant Alzheimer’s Disease: A Review and Proposal for the Prevention of Alzheimer’s Disease. Alzheimers Res. Ther. 2011, 3 (1), 1. https://doi.org/10.1186/alzrt59.

(24) Korenberg, J. R.; Pulst, S.-M.; Neve, R. L.; West, R. The Alzheimer Amyloid Precursor Protein Maps to Human Chromosome 21 Bands Q21.105–Q21.05. Genomics 1989, 5 (1), 124–127. https://doi.org/10.1016/0888-7543(89)90095-5.

(25) Mann, D. M. A.; Esiri, M. M. The Pattern of Acquisition of Plaques and Tangles in the Brains of Patients under 50 Years of Age with Down’s Syndrome. J. Neurol. Sci. 1989, 89 (2), 169–179. https://doi.org/10.1016/0022-510X(89)90019-1.

(26) Castro, P.; Zaman, S.; Holland, A. Alzheimer’s Disease in People with Down’s Syndrome: The Prospects for and the Challenges of Developing Preventative Treatments. J. Neurol. 2017, 264 (4), 804–813. https://doi.org/10.1007/s00415-016-8308-8.

(27) Hof, P. R.; Bouras, C.; Perl, D. P.; Sparks, D. L.; Mehta, N.; Morrison, J. H. Age-Related Distribution of Neuropathologic Changes in the Cerebral Cortex of Patients With Down’s Syndrome: Quantitative Regional Analysis and Comparison With Alzheimer’s Disease. Arch. Neurol. 1995, 52 (4), 379–391. https://doi.org/10.1001/archneur.1995.00540280065020.

(28) Davidson, Y. S.; Robinson, A.; Prasher, V. P.; Mann, D. M. A. The Age of Onset and Evolution of Braak Tangle Stage and Thal Amyloid Pathology of Alzheimer’s Disease in Individuals with Down Syndrome. Acta Neuropathol. Commun. 2018, 6. https://doi.org/10.1186/s40478-018-0559-4.

(29) Ballard, C.; Mobley, W.; Hardy, J.; Williams, G.; Corbett, A. Dementia in Down’s Syndrome. Lancet Neurol. 2016, 15 (6), 622–636. https://doi.org/10.1016/S1474-4422(16)00063-6.

(30) Fortea, J.; Vilaplana, E.; Carmona-Iragui, M.; Benejam, B.; Videla, L.; Barroeta, I.; Fernández, S.; Altuna, M.; Pegueroles, J.; Montal, V.; Valldeneu, S.; Giménez, S.; González-Ortiz, S.; Muñoz, L.; Estellés, T.; Illán-Gala, I.; Belbin, O.; Camacho, V.; Wilson, L. R.; Annus, T.; Osorio, R. S.; Videla, S.; Lehmann, S.; Holland, A. J.; Alcolea, D.; Clarimón, J.; Zaman, S. H.; Blesa, R.; Lleó, A. Clinical and Biomarker Changes of Alzheimer’s Disease in Adults with Down Syndrome: A Cross-Sectional Study. The Lancet 2020, 395 (10242), 1988–1997. https://doi.org/10.1016/S0140-6736(20)30689-9.

(31) Lott, I. T.; Head, E. Dementia in Down Syndrome: Unique Insights for Alzheimer Disease Research. Nat. Rev. Neurol. 2019, 15 (3), 135–147. https://doi.org/10.1038/s41582-018-0132-6.

(32) Snyder, H. M.; Bain, L. J.; Brickman, A. M.; Carrillo, M. C.; Esbensen, A. J.; Espinosa, J. M.; Fernandez, F.; Fortea, J.; Hartley, S. L.; Head, E.; Hendrix, J.; Kishnani, P. S.; Lai, F.; Lao, P.; Lemere, C.; Mobley, W.; Mufson, E. J.; Potter, H.; Zaman, S. H.; Granholm, A.-C.; Rosas, H. D.; Strydom, A.; Whitten, M. S.; Rafii, M. S. Further Understanding the Connection between Alzheimer’s Disease and Down Syndrome. Alzheimers Dement. 2020, 16 (7), 1065–1077. https://doi.org/10.1002/alz.12112.

(33) Abrahamson, E. E.; Head, E.; Lott, I. T.; Handen, B. L.; Mufson, E. J.; Christian, B. T.; Klunk, W. E.; Ikonomovic, M. D. Neuropathological Correlates of Amyloid PET Imaging in Down Syndrome. Dev. Neurobiol. 2019, 79 (7), 750–766. https://doi.org/10.1002/dneu.22713.

(34) Scholl, M.; Wall, A.; Thordardottir, S.; Ferreira, D.; Bogdanovic, N.; Langstrom, B.; Almkvist, O.; Graff, C.; Nordberg, A. Low PiB PET Retention in Presence of Pathologic CSF Biomarkers in Arctic APP Mutation Carriers. Neurology 2012, 79 (3), 229–236. https://doi.org/10.1212/WNL.0b013e31825fdf18.

(35) Rosen, R. F.; Ciliax, B. J.; Wingo, T. S.; Gearing, M.; Dooyema, J.; Lah, J. J.; Ghiso, J. A.; LeVine, H.; Walker, L. C. Deficient High-Affinity Binding of Pittsburgh Compound B in a Case of Alzheimer’s Disease. Acta Neuropathol. (Berl.) 2010, 119 (2), 221–233. https://doi.org/10.1007/s00401-009-0583-3.

(36) Matveev, S. V.; Spielmann, H. P.; Metts, B. M.; Chen, J.; Onono, F.; Zhu, H.; Scheff, S. W.; Walker, L. C.; LeVine, H. A Distinct Subfraction of Aβ Is Responsible for the High-Affinity Pittsburgh Compound B-Binding Site in Alzheimer’s Disease Brain. J. Neurochem. 2014, 131 (3), 356–368. https://doi.org/10.1111/jnc.12815.

(37) Frid, P.; Anisimov, S. V.; Popovic, N. Congo Red and Protein Aggregation in Neurodegenerative Diseases. Brain Res. Rev. 2007, 53 (1), 135–160. https://doi.org/10.1016/j.brainresrev.2006.08.001.

(38) Vassar, P. S.; Culling, C. F. Fluorescent Stains, with Special Reference to Amyloid and Connective Tissues. Arch. Pathol. 1959, 68, 487–498.

(39) Jun, Y. W.; Cho, S. W.; Jung, J.; Huh, Y.; Kim, Y.; Kim, D.; Ahn, K. H. Frontiers in Probing Alzheimer’s Disease Biomarkers with Fluorescent Small Molecules. ACS Cent. Sci. 2019, 5 (2), 209–217. https://doi.org/10.1021/acscentsci.8b00951.

(40) Stöhr, J.; Wu, H.; Nick, M.; Wu, Y.; Bhate, M.; Condello, C.; Johnson, N.; Rodgers, J.; Lemmin, T.; Acharya, S.; Becker, J.; Robinson, K.; Kelly, M. J. S.; Gai, F.; Stubbs, G.; Prusiner, S. B.; DeGrado, W. F. A 31-Residue Peptide Induces Aggregation of Tau’s Microtubule-Binding Region in Cells. Nat. Chem. 2017, 9 (9), 874–881. https://doi.org/10.1038/nchem.2754.

(41) Belloy, M. E.; Napolioni, V.; Greicius, M. D. A Quarter Century of APOE and Alzheimer’s Disease: Progress to Date and the Path Forward. Neuron 2019, 101 (5), 820–838. https://doi.org/10.1016/j.neuron.2019.01.056.

(42) Prasher, V. P.; Sajith, S. G.; Rees, S. D.; Patel, A.; Tewari, S.; Schupf, N.; Zigman, W. B. Significant Effect of APOE Epsilon 4 Genotype on the Risk of Dementia in Alzheimer’s Disease and Mortality in Persons with Down Syndrome. Int. J. Geriatr. Psychiatry 2008, 23 (11), 1134–1140. https://doi.org/10.1002/gps.2039.

(43) Mayeux, R.; Stern, Y.; Ottman, R.; Tatemichi, T. K.; Tang, M. X.; Maestre, G.; Ngai, C.; Tycko, B.; Ginsberg, H. The Apolipoprotein Epsilon 4 Allele in Patients with Alzheimer’s Disease. Ann. Neurol. 1993, 34 (5), 752–754. https://doi.org/10.1002/ana.410340527.

(44) Ossenkoppele, R.; Smith, R.; Mattsson-Carlgren, N.; Groot, C.; Leuzy, A.; Strandberg, O.; Palmqvist, S.; Olsson, T.; Jögi, J.; Stormrud, E.; Cho, H.; Ryu, Y. H.; Choi, J. Y.; Boxer, A. L.; Gorno-Tempini, M. L.; Miller, B. L.; Soleimani-Meigooni, D.; Iaccarino, L.; La Joie, R.; Baker, S.; Borroni, E.; Klein, G.; Pontecorvo, M. J.; Devous, M. D.; Jagust, W. J.; Lyoo, C. H.; Rabinovici, G. D.; Hansson, O. Accuracy of Tau Positron Emission Tomography as a Prognostic Marker in Preclinical and Prodromal Alzheimer Disease: A Head-to-Head Comparison Against Amyloid Positron Emission Tomography and Magnetic Resonance Imaging. JAMA Neurol. 2021. https://doi.org/10.1001/jamaneurol.2021.1858.

(45) Aoyagi, A.; Condello, C.; Stöhr, J.; Yue, W.; Rivera, B. M.; Lee, J. C.; Woerman, A. L.; Halliday, G.; van Duinen, S.; Ingelsson, M.; Lannfelt, L.; Graff, C.; Bird, T. D.; Keene, C. D.; Seeley, W. W.; DeGrado, W. F.; Prusiner, S. B. Aβ and Tau Prion-like Activities Decline with Longevity in the Alzheimer’s Disease Human Brain. Sci. Transl. Med. 2019, 11 (490). https://doi.org/10.1126/scitranslmed.aat8462.

(46) Leverenz, J. B.; Raskind, M. A. Early Amyloid Deposition in the Medial Temporal Lobe of Young Down Syndrome Patients: A Regional Quantitative Analysis. Exp. Neurol. 1998, 150 (2), 296–304. https://doi.org/10.1006/exnr.1997.6777.

(47) Lemere, C. A.; Blusztajn, J. K.; Yamaguchi, H.; Wisniewski, T.; Saido, T. C.; Selkoe, D. J. Sequence of Deposition of Heterogeneous Amyloid β-Peptides and APO E in Down Syndrome: Implications for Initial Events in Amyloid Plaque Formation. Neurobiol. Dis. 1996, 3 (1), 16–32.

(48) Iwatsubo, T.; Mann, D. M. A.; Odaka, A.; Suzuki, N.; Ihara, Y. Amyloid β Protein (Aβ) Deposition: Aβ42(43) Precedes Aβ40 in down Syndrome. Ann. Neurol. 1995, 37 (3), 294–299. https://doi.org/10.1002/ana.410370305.

(49) Wesseling, H.; Mair, W.; Kumar, M.; Schlaffner, C. N.; Tang, S.; Beerepoot, P.; Fatou, B.; Guise, A. J.; Cheng, L.; Takeda, S.; Muntel, J.; Rotunno, M. S.; Dujardin, S.; Davies, P.; Kosik, K. S.; Miller, B. L.; Berretta, S.; Hedreen, J. C.; Grinberg, L. T.; Seeley, W. W.; Hyman, B. T.; Steen, H.; Steen, J. A. Tau PTM Profiles Identify Patient Heterogeneity and Stages of Alzheimer’s Disease. Cell 2020, 183 (6), 1699–1713.e13. https://doi.org/10.1016/j.cell.2020.10.029.

(50) Wisniewski, K. E.; Wisniewski, H. M.; Wen, G. Y. Occurrence of Neuropathological Changes and Dementia of Alzheimer’s Disease in Down’s Syndrome. Ann. Neurol. 1985, 17 (3), 278–282. https://doi.org/10.1002/ana.410170310.

(51) Braak, H.; Braak, E. Neuropathological Stageing of Alzheimer-Related Changes. Acta Neuropathol. (Berl.) 1991, 82 (4), 239–259. https://doi.org/10.1007/BF00308809.

(52) van Dyck, C. H. Anti-Amyloid-β Monoclonal Antibodies for Alzheimer’s Disease: Pitfalls and Promise. Biol. Psychiatry 2018, 83 (4), 311–319. https://doi.org/10.1016/j.biopsych.2017.08.010.

(53) Armstrong, R. A. The Molecular Biology of Senile Plaques and Neurofibrillary Tangles in Alzheimer’s Disease. Folia Neuropathol. 2009, 47 (4), 289–299.

(54) Harada, R.; Okamura, N.; Furumoto, S.; Yoshikawa, T.; Arai, H.; Yanai, K.; Kudo, Y. Use of a Benzimidazole Derivative BF-188 in Fluorescence Multispectral Imaging for Selective Visualization of Tau Protein Fibrils in the Alzheimer’s Disease Brain. Mol. Imaging Biol. 2014, 16 (1), 19–27. https://doi.org/10.1007/s11307-013-0667-2.

(55) Simon, R. A.; Shirani, H.; Åslund, K. O. A.; Bäck, M.; Haroutunian, V.; Gandy, S.; Nilsson, K. P. R. Pentameric Thiophene-Based Ligands That Spectrally Discriminate Amyloid-β and Tau Aggregates Display Distinct Solvatochromism and Viscosity-Induced Spectral Shifts. Chem. - Eur. J. 2014, 20 (39), 12537–12543. https://doi.org/10.1002/chem.201402890.

(56) Condello, C.; Yuan, P.; Schain, A.; Grutzendler, J. Microglia Constitute a Barrier That Prevents Neurotoxic Protofibrillar Aβ42 Hotspots around Plaques. Nat. Commun. 2015, 6, 6176. https://doi.org/10.1038/ncomms7176.

(57) Ulrich, J. D.; Ulland, T. K.; Mahan, T. E.; Nyström, S.; Nilsson, K. P.; Song, W. M.; Zhou, Y.; Reinartz, M.; Choi, S.; Jiang, H.; Stewart, F. R.; Anderson, E.; Wang, Y.; Colonna, M.; Holtzman, D. M. ApoE Facilitates the Microglial Response to Amyloid Plaque Pathology. J. Exp. Med. 2018, 215 (4), 1047–1058. https://doi.org/10.1084/jem.20171265.

(58) Yuan, P.; Condello, C.; Keene, C. D.; Wang, Y.; Bird, T. D.; Paul, S. M.; Luo, W.; Colonna, M.; Baddeley, D.; Grutzendler, J. TREM2 Haplodeficiency in Mice and Humans Impairs the Microglia Barrier Function Leading to Decreased Amyloid Compaction and Severe Axonal Dystrophy. Neuron 2016, 90 (4), 724–739. https://doi.org/10.1016/j.neuron.2016.05.003.

(59) Xue, Q.-S.; Streit, W. J. Microglial Pathology in Down Syndrome. Acta Neuropathol. (Berl.) 2011, 122 (4), 455–466. https://doi.org/10.1007/s00401-011-0864-5.

(60) Martini, A. C.; Helman, A. M.; McCarty, K. L.; Lott, I. T.; Doran, E.; Schmitt, F. A.; Head, E. Distribution of Microglial Phenotypes as a Function of Age and Alzheimer’s Disease Neuropathology in the Brains of People with Down Syndrome. Alzheimers Dement. Amst. Neth. 2020, 12 (1), e12113. https://doi.org/10.1002/dad2.12113.

(61) Ovchinnikov, D. A.; Korn, O.; Virshup, I.; Wells, C. A.; Wolvetang, E. J. The Impact of APP on Alzheimer-like Pathogenesis and Gene Expression in Down Syndrome IPSC-Derived Neurons. Stem Cell Rep. 2018, 11 (1), 32–42. https://doi.org/10.1016/j.stemcr.2018.05.004.

(62) Doran, E.; Keator, D.; Head, E.; Phelan, M. J.; Kim, R.; Totoiu, M.; Barrio, J. R.; Small, G. W.; Potkin, S. G.; Lott, I. T. Down Syndrome, Partial Trisomy 21, and Absence of Alzheimer’s Disease: The Role of APP. J. Alzheimers Dis. 2017, 56 (2), 459–470. https://doi.org/10.3233/JAD-160836.

(63) Wiseman, F. K.; Pulford, L. J.; Barkus, C.; Liao, F.; Portelius, E.; Webb, R.; Chávez-Gutiérrez, L.; Cleverley, K.; Noy, S.; Sheppard, O.; Collins, T.; Powell, C.; Sarell, C. J.; Rickman, M.; Choong, X.; Tosh, J. L.; Siganporia, C.; Whittaker, H. T.; Stewart, F.; Szaruga, M.; Murphy, M. P.; Blennow, K.; de Strooper, B.; Zetterberg, H.; Bannerman, D.; Holtzman, D. M.; Tybulewicz, V. L. J.; Fisher, E. M. C. Trisomy of Human Chromosome 21 Enhances Amyloid-β Deposition Independently of an Extra Copy of APP. Brain 2018, 141 (8), 2457–2474. https://doi.org/10.1093/brain/awy159.

(64) Ryoo, S.-R.; Jeong, H. K.; Radnaabazar, C.; Yoo, J.-J.; Cho, H.-J.; Lee, H.-W.; Kim, I.-S.; Cheon, Y.-H.; Ahn, Y. S.; Chung, S.-H.; Song, W.-J. DYRK1A-Mediated Hyperphosphorylation of Tau. J. Biol. Chem. 2007, 282 (48), 34850–34857. https://doi.org/10.1074/jbc.M707358200.

(65) Wegiel, J.; Dowjat, K.; Kaczmarski, W.; Kuchna, I.; Nowicki, K.; Frackowiak, J.; Mazur Kolecka, B.; Wegiel, J.; Silverman, W. P.; Reisberg, B.; deLeon, M.; Wisniewski, T.; Gong, C.-X.; Liu, F.; Adayev, T.; Chen-Hwang, M.-C.; Hwang, Y.-W. The Role of Overexpressed DYRK1A Protein in the Early Onset of Neurofibrillary Degeneration in Down Syndrome. Acta Neuropathol. (Berl.) 2008, 116 (4), 391–407. https://doi.org/10.1007/s00401-008-0419-6.

(66) Song, W.-J.; Song, E.-A. C.; Choi, S.-H.; Baik, H.-H.; Jin, B. K.; Kim, J. H.; Chung, S.-H. Dyrk1A-Mediated Phosphorylation of RCAN1 Promotes the Formation of Insoluble RCAN1 Aggregates. Neurosci. Lett. 2013, 554, 135–140. https://doi.org/10.1016/j.neulet.2013.08.066.

(67) Giles, K.; Berry, D. B.; Condello, C.; Hawley, R. C.; Gallardo-Godoy, A.; Bryant, C.; Oehler, A.; Elepano, M.; Bhardwaj, S.; Patel, S.; Silber, B. M.; Guan, S.; DeArmond, S. J.; Renslo, A. R.; Prusiner, S. B. Different 2-Aminothiazole Therapeutics Produce Distinct Patterns of Scrapie Prion Neuropathology in Mouse Brains. J. Pharmacol. Exp. Ther. 2015, 355 (1), 2–12. https://doi.org/10.1124/jpet.115.224659.

(68) Berry, D. B.; Lu, D.; Geva, M.; Watts, J. C.; Bhardwaj, S.; Oehler, A.; Renslo, A. R.; DeArmond, S. J.; Prusiner, S. B.; Giles, K. Drug Resistance Confounding Prion Therapeutics. Proc. Natl. Acad. Sci. 2013, 110 (44), E4160–E4169. https://doi.org/10.1073/pnas.1317164110.

(69) Selkoe, D. J. Alzheimer Disease and Aducanumab: Adjusting Our Approach. Nat. Rev. Neurol. 2019, 15 (7), 365–366. https://doi.org/10.1038/s41582-019-0205-1.

(70) Armstrong, R. A.; Smith, C. U. M. β-Amyloid (β/A4) Deposition in the Medial Temporal Lobe in Down’s Syndrome: Effects of Brain Region and Patient Age. Neurobiol. Dis. 1994, 1 (3), 139–144. https://doi.org/10.1006/nbdi.1994.0017.

(71) Annus, T.; Wilson, L. R.; Hong, Y. T.; Acosta–Cabronero, J.; Fryer, T. D.; Cardenas–Blanco, A.; Smith, R.; Boros, I.; Coles, J. P.; Aigbirhio, F. I.; Menon, D. K.; Zaman, S. H.; Nestor, P. J.; Holland, A. J. The Pattern of Amyloid Accumulation in the Brains of Adults with Down Syndrome. Alzheimers Dement. 2016, 12 (5), 538–545. https://doi.org/10.1016/j.jalz.2015.07.490.

(72) Zhong, L.; Xie, Y.-Z.; Cao, T.-T.; Wang, Z.; Wang, T.; Li, X.; Shen, R.-C.; Xu, H.; Bu, G.; Chen, X.-F. A Rapid and Cost-Effective Method for Genotyping Apolipoprotein E Gene Polymorphism. Mol. Neurodegener. 2016, 11 (1), 2. https://doi.org/10.1186/s13024-016-0069-4.

(73) MATLAB; The MathWorks Inc.: Natick, Massachusetts, 2021.

